# Identification and characterization of constrained non-exonic bases lacking predictive epigenomic and transcription factor binding annotations

**DOI:** 10.1101/722876

**Authors:** Olivera Grujic, Tanya N. Phung, Soo Bin Kwon, Adriana Arneson, Yuju Lee, Kirk E. Lohmueller, Jason Ernst

## Abstract

Annotations of evolutionarily constraint provide important information for variant prioritization. Genome-wide maps of epigenomic marks and transcription factor binding provide complementary information for interpreting a subset of such prioritized variants. Here we developed the Constrained Non-Exonic Predictor (CNEP) to quantify the evidence of each base in the human genome being in a constrained non-exonic element from over 60,000 epigenomic and transcription factor binding features. We find that the CNEP score outperforms baseline and related existing scores at predicting constrained non-exonic bases from such data. However, a subset of such bases are still not well predicted by CNEP. We developed a complementary Conservation Signature Score by CNEP (CSS-CNEP) using conservation state and constrained element annotations that is predictive of those bases. Using human genetic variation, regulatory sequence motifs, mouse epigenomic data, and retrospectively considered additional human data we further characterize the nature of constrained non-exonic bases with low CNEP scores.

## Introduction

A large majority of genetic variation associated with common disease falls into non-exonic regions of the human genome^1^. Annotations of evolutionarily constrained elements have either been used directly or as an important feature to integrative methods for prioritizing potentially deleterious non-exonic mutations^2–8^. Supporting the importance of these annotations, heritability analyses have suggested they are heavily enriched for disease associated variants^9^.

When studying a non-coding variant prioritized based on comparative genomic information, the problem of determining the cell or tissue type of activity and mechanism by which such a variant exerts influence remains. Genome-wide maps of histone modifications and variants, open chromatin, chromatin state annotations, and transcription factor (TF) binding can give insights^10–14^. However, such data is specific to the condition and cell or tissue type in which the experiments underlying them were conducted. Previous analyses have shown that while there is an enrichment for evolutionarily constrained bases in epigenomic or transcription factor binding annotations of regulatory activity, some evolutionary constrained bases lack informative annotations^4,12,15–20^.

When investigating the role of a prioritized variant in a constrained non-exonic element with a compendium of epigenomic and TF binding, an initial question is whether the constraint can even be explained by information in the compendium of data. However, with tens of thousands epigenomic and TF binding data sets available, answering this question is not straightforward. Integrative scores such as CADD^7^ that combine epigenomic data and TF binding features with conservation features cannot be directly applied to answer such a question, since a base could receive a high score based on the conservation features even if informative epigenomic or TF binding annotations were lacking.

A few scores have been proposed that can be used to quantify information in epigenomic or TF binding data. The FitCons and FitCons2 methods^21,22^ quantified information in epigenomic data with polymorphism and divergence information to provide cell type specific estimates of fitness. A ‘conservation-associated activity score’ was recently introduced, which provided a cell type specific score of Segway annotations based on the genome-wide distribution of an evolutionary constraint score in the annotation in the cell type^23^. Additionally, for both of these approaches a single summary score of the cell type specific scores was also computed. While informative, if one is interested in a single score summarizing information in a compendium of epigenomic and TF data, an approach that defines the single score without going through cell type specific scores would have greater flexibility in terms of what data sets are used to and how information within them are combined to produce the score.

Here we developed a method, the Constrained Non-Exonic Predictor (CNEP), designed to produce a single score, without respect to cell type, that reflects the probability that a base will be in an evolutionarily constrained non-exonic element from information in a large-scale compendium of epigenomic and TF binding data. We focus specifically on non-exonic bases since they comprise a much larger portion of the genome and are less well annotated compared to exons. Also, constraint in such bases is expected to be largely associated with distinct patterns of epigenomic marks and TF binding relative to that found in exons.

We applied CNEP to provide a score for each base of the human genome, summarizing the probability of a base being in a constrained non-exonic element based on more than 60,000 features defined from a large compendium of epigenomic and TF binding data. We show that the CNEP score outperforms baseline and related existing scores at predicting such bases. While the CNEP score is able to effectively predict many bases in constrained non-exonic elements, a substantial portion of bases were still not well predicted by CNEP.

We also developed a complementary Conservation Signature Score by CNEP (CSS-CNEP), which can be used to differentiate bases in constrained non-exonic elements with low CNEP scores that are more likely due to being false calls of constraint from those active in cell types and conditions not represented in the compendium. The CSS-CNEP computes the expected CNEP score at a position based on the recently developed ConsHMM conservation state annotations^24^, thus leveraging detailed alignment pattern information, along with constrained element annotations. We show that when considering bases in constrained non-exonic elements, that CSS-CNEP is partially predictive of the subset of those bases that received a low CNEP score.

Using human genetic variation data, we show that a portion of the constrained non-exonic bases that were not well predicted by CNEP truly appear to be under constraint. We conducted additional analyses using regulatory sequence motif annotations, chromatin accessibility data from mouse, and retrospectively considered additional human epigenomic and TF binding data to provide insights into the potential role of constrained non-exonic bases not captured by commonly used compendia of epigenomic and TF binding data. We also analyzed bases that receive a high CNEP score, but are not in a constrained element, a portion of which may correspond to adaptive evolution or changing selective effects over time. CNEP and CSS-CNEP are resources for analyzing the genome and variants in terms of large-scale epigenomic and TF binding data, evolutionary constraint annotations, and their relationship.

## Results

### Constrained Non-Exonic Predictor method

We developed the Constrained Non-Exonic Predictor (CNEP) to make a probabilistic prediction based on features defined from large-scale epigenomics and TF binding data as to whether a base in the human genome will be in a constrained non-exonic element previously called from comparative genomics sequence analysis (**Fig. 1**). We applied CNEP with 63,741 features derived from overlap of peak calls in experiments mapping TF binding, histone modifications, and chromatin accessibility, as well as TF binding footprint calls from Digital Genomic Footprinting, and chromatin state annotations from ChromHMM (**Supplementary Table 1-2; Methods**). For simplicity of presentation we refer to TF binding to also include general factor binding that is not necessarily sequence specific. The features are based on data produced by the Roadmap Epigenomics and ENCODE consortia, or previously curated and reprocessed public data from the ReMap or ChIP-atlas databases^11,14,25–27^. ReMap provides a uniform processing TF binding data from the Gene Expression Omnibus and ArrayExpress, while ChIP-atlas provides a uniform processing of Sequence Read Archive data for epigenomic and TF binding data^26,28^.

**Figure 1:**
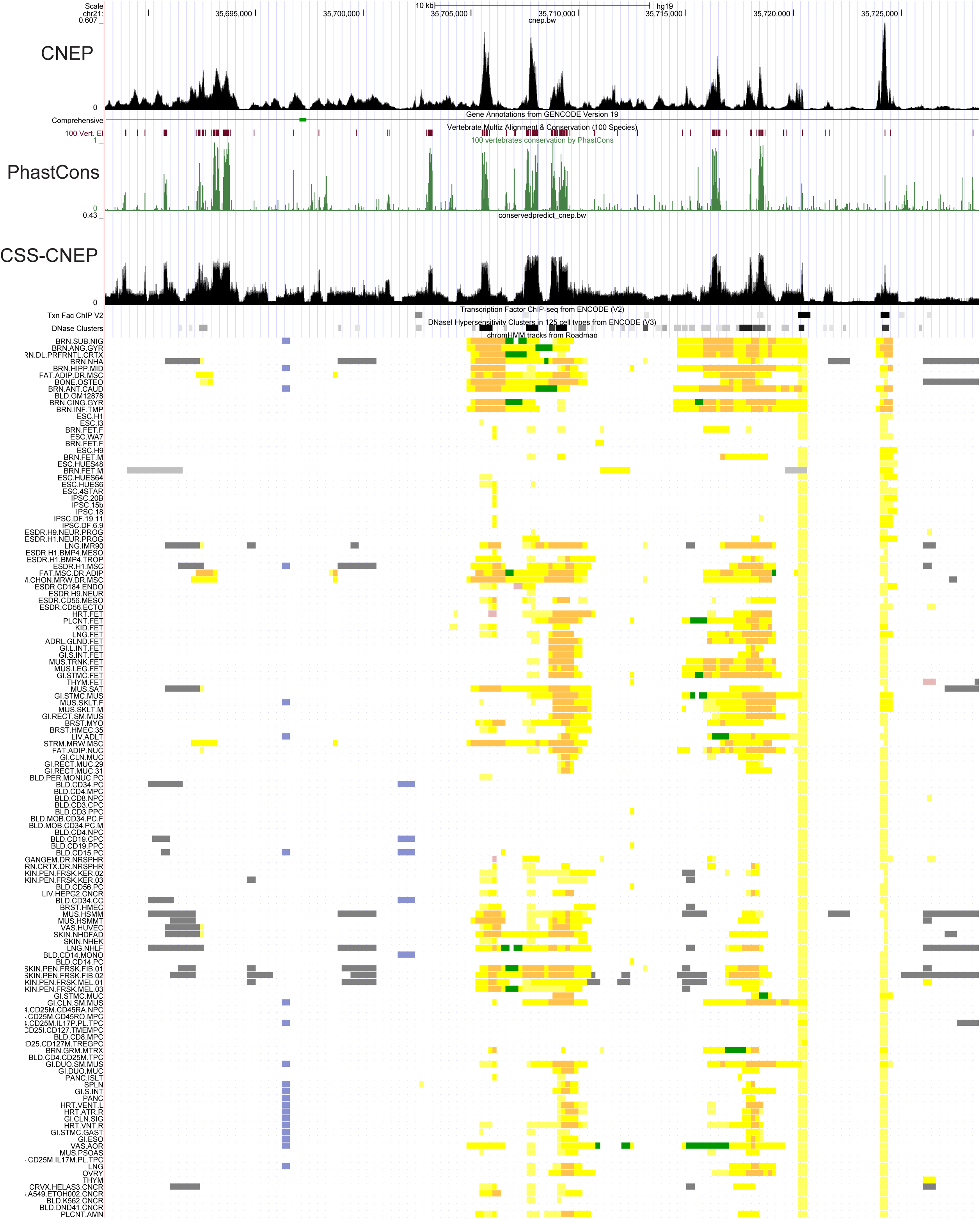
Example of Constrained Non-exonic Predictor (CNEP) scores. An example genomic locus illustrating CNEP scores. The top line is the CNEP score track. In general, the CNEP score ranges between 0 and 1, but in this image the y-scale is capped at 0.5. Below the CNEP score track is the GENCODE gene annotation track, PhastCons element track, and then the PhastCons score track. Below the PhastCons score track is the Conservation Signature Score by CNEP (CSS-CNEP), followed by the UCSC Genome Browser ENCODE TF binding summary track (Txn Fac ChIP V2), and the ENCODE DNase I summary track (DNase Clusters V3). These tracks are then followed by chromatin state annotation across 127 samples based on a previously defined 25-state ChromHMM annotation based on imputed data^36^. A color legend for the chromatin state annotations is also available in **Fig. 2d**.

CNEP trains an ensemble of logistic regression classifiers to discriminate between bases overlapping evolutionarily constrained elements outside of a GENCODE annotated exon and those bases in the rest of the genome (**Methods**). While constrained element annotations are used to provide labels for training, we emphasize that CNEP does not use any features based on comparative genomics data for predictions. We used an ensemble of logistic regression classifiers as they provide a probabilistic output and are robust. For each chromosome, CNEP trains a separate set of classifiers based on subsamples of positions from all chromosomes except the target chromosome. CNEP then makes a probabilistic prediction between 0 and 1 for each base on the target chromosome as to whether the base is in a constrained non-exonic element. We applied CNEP using for labels constrained element sets previously produced by four different methods: PhastCons^5,29^, GERP++^2^, SiPhy-pi and SiPhy-omega^3,4^ (**Methods**).

Predictions based on training on any two different constrained element sets were all highly correlated, with correlations ranging from 0.88 to 0.93 (**Supplementary Fig. 1**). To reduce biases associated with the choice of one specific constrained element set and to simplify downstream analyses, we averaged the predictions to derive a single score, which we termed the CNEP score (**Fig. 2a, Supplementary Fig. 2**). We confirmed that the CNEP score, based on averaging the four predictions, was even more highly correlated with scores based on a single constrained element set with correlations ranging between 0.96 and 0.98 (**Supplementary Fig. 1**).

**Figure 2:**
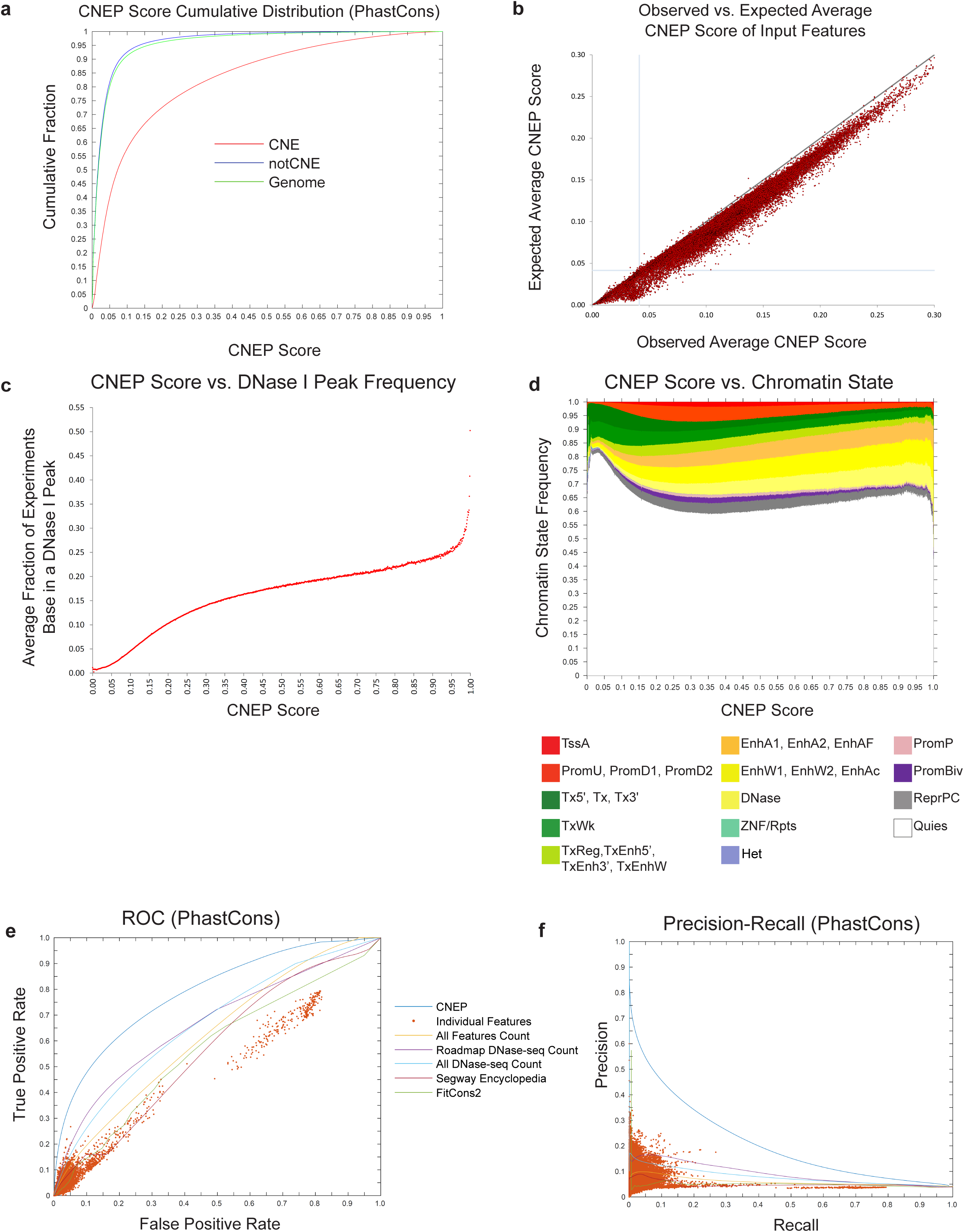
Properties of the CNEP score. **(a)** The graph shows the cumulative distribution of the CNEP score genome-wide (green), in PhastCons constrained non-exonic (CNE) bases (red), and bases that are not in PhastCons constrained elements and also not in exons (notCNE) (blue). **(b)** A scatter plot with each point corresponding to one feature that CNEP uses. The x-axis shows the average CNEP score in bases that have the feature present, while the y-axis shows the expected CNEP score based on the feature’s overlap with constrained non-exonic bases. Only 48,364 features that cover at least 200kb are shown. The full table corresponding to these values can be found in **Supplementary Table 2**. The diagonal line is the y=x line. The vertical line corresponds to the genome-wide average CNEP score. The horizontal line corresponds to the genome-wide expected average CNEP score. **(c)** A plot showing the average fraction of the 350 Roadmap DNase I experiments in which the base overlaps a called peak for each CNEP score value, rounded to the nearest 0.001, covering at least 1000 bases. In total, there was 1000 such values. **(d)** A plot showing the average fraction of bases assigned across the 127 epigenomes to each of 14-groups based on 25 ChromHMM chromatin states previously assigned the same color^36^ for each CNEP score value, rounded to the nearest 0.001. A color with the state abbreviations is displayed at the bottom of the panel. **(e)** A plot of the ROC curve for the CNEP score predicting PhastCons non-exonic bases. The area under this curve is 0.79. Also shown is the performance of individual features and several baseline or existing scores (**Methods**). **(f)** A plot of the precision-recall curve for the CNEP score identifying PhastCons non-exonic bases and the same individual features and baseline and existing scores as (e). ROC and precision-recall curves for other constrained element sets can be found in **Supplementary Fig. 3**.

### CNEP score associates with signatures of regulatory activity

We next investigated the relationship between CNEP scores and the input features to CNEP. For each input feature, we computed the observed genome-wide average CNEP score of bases overlapped by the feature and compared it to the expected average CNEP score computed based on the proportion of the feature’s bases overlapping with constrained non-exonic elements on average for the four element sets (**Methods, Supplementary Table 2, Fig. 2b**). We found that the observed and expected average CNEP score showed strong agreement with a pearson correlation of 0.99 for those features covering at least 200kb (**Fig. 2b**). We also confirmed that the genome-wide actual observed average CNEP score of 0.0419 was close to the expected average score of 0.0415 based on the average genome coverage of the four constrained element sets.

We next investigated whether bases that received a higher CNEP score were more likely to show signatures of regulatory activity in more experiments or cell and tissue types that are subsets of input features to CNEP. Specifically, we first analyzed a set of 350 DNase I hypersensitivity experiments from the Roadmap Epigenomics project. We note that, 51% of bases in the genome were covered by at least one peak highlighting the limited specificity in the annotation of whether a base in the genome overlaps a DNase I peak in any experiment. We computed the average number of these experiments in which a base would be covered by a peak as a function of the CNEP score (**Fig. 2c**). On average, a base in the genome was covered in 1.9% experiments. Bases that received a higher CNEP score tended to be in a DNase I hypersensitivity peak in more experiments. For example, bases with a CNEP score of 0.500 were in a peak in 18.0% of the experiments on average. We saw a similar pattern when considering the chromatin state frequency from a chromatin state model defined across 127 cell and tissue types as a function of the CNEP score, which showed on average a greater presence of candidate enhancer or promoter chromatin states for greater CNEP scores (**Fig. 2d**).

### CNEP outperforms baselines and related scores at predicting constrained non-exonic elements

We next analyzed the extent to which the CNEP score is able to predict bases in constrained non-exonic elements out of all other bases in the genome using Receiving Operator Characteristic (ROC) curves and precision-recall curves (**Fig. 2e,f**, **Supplementary Fig. 3**). We obtained area under the ROC curves in the range 0.79 to 0.86 depending on the constrained element set being predicted, with the area under the curve (AUC) values for PhastCons elements being lower than the other three element sets. The lower AUC for PhastCons might be related at least in part to this being the only element set of the three defined using alignment information that included non-mammalian vertebrates, which can make it more vulnerable to possible alignment errors, or due to the higher resolution at which the method calls constrained elements^24^.

**Figure 3:**
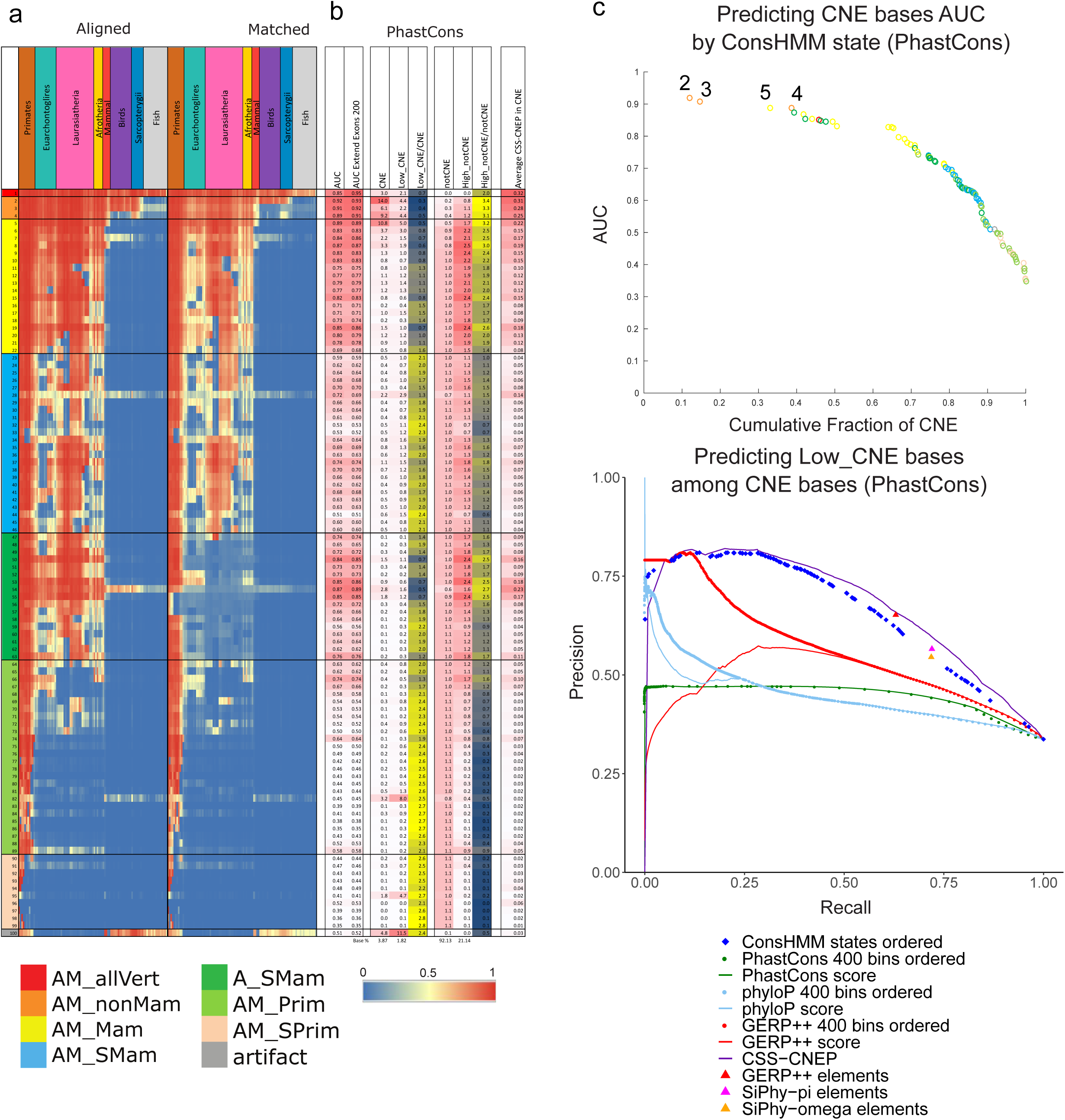
CNE prediction depends on conservation state. **(a)** Heatmap representation of conservation state parameters of the ConsHMM conservation state model defined in Ref. 24. Rows correspond to different conservation states. The states were previously clustered into eight groups based on these parameters and colored accordingly. The left half indicates for each state the probability of each species having a nucleotide aligning the human reference genome, regardless of whether it matches the human reference. The right half indicates for each state the probability of each species having a nucleotide matching the human reference genome. Individual columns correspond to species, the names of which are available in Ref. 24. The major groups of species are colored and labeled. Color scale for the heatmap is shown at the bottom. **(b)** The first column reports the genome % of each state excluding chrY. The second column contains the AUC of the CNEP score for predicting CNE bases in each state, where for this and the remaining columns the constrained elements are from PhastCons. CNE bases that are not in the target conservation state are excluded when computing the AUC. The next column reports the AUC when exons are first extended by 200bp. The next three columns contain the fold enrichment for CNE bases, Low_CNE bases, and the ratio of the enrichment of Low_CNE bases to CNE bases. The next three columns contain the fold enrichment for notCNE bases, High_notCNE bases, and the ratio of the enrichment of High_notCNE bases to notCNE bases. The last column shows the average CSS-CNEP score in CNE bases in the state. Adjacent pairs of columns on a red-white color scale are on the same color scale. The other columns are on a column specific color scale. The bottom row gives the base % of the genome for the four sets. Results based on all the constrained element sets can be found in **Supplementary Fig. 6. (c)** Plot showing the AUC values for each ROC curve for predicting PhastCons CNE bases in specific ConsHMM conservation states shown in **Supplementary Fig. 7a**. The AUC values are displayed from left to right in decreasing value and positioned along the x-axis based on the cumulative fraction of PhastCons CNE bases they cover. The points are color-coded based on the conservation state coloring shown in (a). States with the highest AUC values are labeled. Similar plots, but for additional constrained element sets can be found in **Supplementary Fig. 7** and based on excluding bases within 200bp of exons from the positives can be found in **Supplementary Fig. 8. (d)** Precision-recall analysis for predicting PhastCons Low_CNE bases among CNE bases using additional comparative genomics information. In this analysis, Low_CNE bases are positive bases and High_CNE bases are negative bases. The predictions based on the CSS-CNEP score as well as the PhastCons, PhyloP, and GERP++ constraint scores are shown based on ranking from lowest to highest value. Also shown for the PhastCons, PhyloP, and GERP++ scores are precision-recall curves, based on dividing a score into four hundred bins and ordering the bins based on increasing enrichment on a training set containing separate positions than used for the evaluation (**Methods**). The plot also shows the cumulative precision recall of the conservation states when ordered based on enrichment for Low_CNE bases in the training data. Additionally, a single point is shown for each of the other three constrained element sets corresponding to predictions based on bases not covered by them. Similar plots for additional constrained elements can be found in **Supplementary Fig. 9**.

To place the predictive performance of the CNEP score in perspective, we compared to several different baselines and existing scores (**Fig. 2ef, Supplementary Fig. 3**). We first verified that the CNEP score had a better true positive rate at the same false positive rate than any individual input feature and likewise for precision at the same recall. We then compared to a baseline score where we counted the number of input features overlapping a base. We also compared to two additional baseline scores restricted to counting overlapping input features that were for any DNase I hypersensitive experiment or specifically from one of the 350 from Roadmap Epigenomics considered above. In all cases the CNEP score outperformed these baselines. The AUCs were in the range [0.62,0.69] when counting all features and in the range [0.67,0.74] for both DNase I baselines, compared to [0.79,0.86] for CNEP, with the specific value in the range depending on the specific constrained element set. Two related existing scores, Segway Encyclopedia ‘conservation-associated activity score’^23^ and FitCons2 ‘cell-type integrated scores’^21^ had even lower AUC values, in the ranges [0.57,0.66] and [0.57,0.62], respectively. We saw similar results when we excluded any base overlapping an exon from the negative set, and when we used a version of the CNEP score based on just the element set being evaluated before averaging scores for different element sets (**Supplementary Table 3**).

### Subset of bases in constrained non-exonic elements are not well predicted by CNEP

We next sought to define and better understand CNEP predictions that disagreed with the annotation of whether a base was in a constrained non-exonic element. Specifically, we defined six sets of bases for each constrained element set. We defined a subset CNE (constrained-non exonic) of bases covered by a constrained element and another subset notCNE as bases not in CNE and not in an exon. We further partitioned the CNE bases into Low_CNE and High_CNE bases depending on whether the CNEP score was below or above the genome-wide average, respectively, and similarly partitioned notCNE bases into Low_notCNE and High_notCNE bases (**Fig. 2a, Supplementary Fig. 2, Supplementary Table 4**). From these definitions, 0.6-1.3% of bases in the genome were in the Low_CNE set, with the specific value in the range depending on the specific constrained element set being considered. This range corresponds to 20.9-34.6% of CNE bases falling into the Low_CNE set, indicating that a substantial fraction of CNE bases called based on sequence constraint had a low CNEP score. For comparison, 19.6-21.3% of the bases in the genome were in the High_notCNE set. These bases received a higher CNEP score than all Low_CNE bases, despite the former not overlapping a base in a constrained element.

While limitations in the resolution of the epigenomic and TF binding data relative to the resolution at which CNE are bases are defined can explain many High_notCNE bases, these limitations cannot explain a substantial portion of High_notCNE bases. Specifically, we computed the prevalence and enrichment of High_notCNE bases as a function of distance to the nearest CNE (**Supplementary Fig. 5**). For example, for PhastCons High_notCNE bases, we found a 2.7-fold enrichment immediately next to a CNE, which decreased as the distance increased. In total 57% of High_notCNE bases were within 200 base pairs of a CNE base, corresponding to a 1.8-fold enrichment. However, 43% of High_notCNE bases were more than 200bp from a CNE base and still received higher CNEP scores than Low_CNE bases.

We also verified proximity to exons provides a limited explanation of Low_CNE bases. Specifically, we computed the prevalence and enrichment of Low_CNE bases as a function of distance to nearest exon (**Supplementary Fig. 6**). For example, for PhastCons CNE bases, we found a 1.8-fold enrichment for bases immediately next to an exon, with the enrichment decreasing further away from the exon. The cumulative enrichment at 200bp away from the nearest exon was 1.3-fold and only contained 10.2% of Low_CNE bases.

### Conservation signatures predictive of CNE bases with a Low CNEP score

A reason that some of the CNE bases might not be predicted by CNEP is that they represent false-positive calls of constraint. We next explored whether there are additional signals present within a multi-species sequence alignment that are predictive of which bases would receive a low CNEP score that were not captured by the original unsupervised model used to call CNE bases. Establishing this, would suggest it is possible to identify CNE bases with a low CNEP score that more likely represent a false call of constraint.

We observed the CNEP scores’ ability to recover CNE bases vary drastically depending on the ConsHMM conservation state^24^ they overlap (**Fig. 3, Supplementary Fig. 6-8**). The ConsHMM conservation states provide single-nucleotide annotations based on the combinatorial and spatial patterns of which species match and align the human genome in a 100-way vertebrate alignment. For example, for PhastCons, while the overall AUC for predicting bases in CNE with CNEP was 0.79, the AUC was as high as 0.92 when considering CNE bases in ConsHMM state 2, which is a state associated with high frequency of all vertebrates except fish aligning to and matching the human reference genome. In contrast, the AUC was less than 0.650 for 56 states, generally having low align frequencies for many mammals, and comprising 17.4% of all CNE bases. After excluding bases within 200bp of exons, the AUC reached as high as 0.95 for state 1. We saw similar trends when considering the enrichment of Low_CNE and High_notCNE bases in each conservation state (**Fig. 3a,b, Supplementary Fig. 6**).

Based on the conservation state assignment and combination of which of the four constrained element sets were present at a base, we predicted the CNEP score for the base, producing what we termed the Conservation Signature Score by CNEP (CSS-CNEP) (**Methods, Fig. 1, 3c, Supplementary Fig. 9**). The CSS-CNEP for a base on one chromosome was computed as the average CNEP score in non-exonic bases that had the same conservation state and combination of constrained elements on other chromosomes. This score was more predictive than any existing constraint score of CNE bases that received a low CNEP score. Additionally, in most cases compared to individual constrained element sets, CSS-CNEP had greater precision at their single recall rate. We also directly analyzed the average CSS-CNEP score in CNE bases for each conservation state, which showed a large variation across conservation states (**Fig. 3, Supplementary Fig. 6**). For example, for PhastCons, CNE bases in states 1 and 2 had average CSS-CNEP scores of 0.32 and 0.30 respectively, while CNE bases in state 100, associated with putative alignment artifacts^24^, had an average CSS-CNEP score of 0.03. The CSS-CNEP thus provides a complementary resource to the CNEP score, allowing one to identify among CNE bases with low CNEP scores those more likely correspond to false positive calls of constraint.

However, the CSS-CNEP is only partially predictive of Low_CNE bases. Consistent with this even for CNE bases in ConsHMM states for which CNEP was most predictive, there was still a substantial subset of bases receiving low CNEP scores. For example, for PhastCons, states 2, 4, and 5 all had greater than four-fold enrichment for Low_CNE bases and contained 19% of Low_CNE bases. Many of these bases may correspond to true calls of constraint, but are not active in experiments provided as input to CNEP.

### Low_CNE bases show evidence for purifying selection within humans

To test whether the set of Low_CNE bases are still enriched for bases under purifying selection in humans despite the limited support from the epigenomics and TF binding data considered, we turned to human genetic variation data. Specifically, we considered a set of 105 unrelated individuals of the Yoruba in Ibadan (YRI) population from the 1000 Genomes Project and first examined the proportional site frequency spectrum (SFS) (**Methods**). Comparing Low_CNE bases to High_notCNE bases, we observed that there is a significant difference in the distribution (p<10^−15^), with a greater proportion of low-frequency variants for Low_CNE bases, especially singletons and doubletons, and a lower proportion of common variants (**Fig. 4a, Supplementary Fig. 10a,c,e**, comparing orange and purple bars). The skew towards low-frequency variants and the deficit in high-frequency variants suggest stronger purifying selection in the Low_CNE bases relative to High_notCNE bases. As an additional evaluation of whether purifying selection has been stronger in the Low_CNE bases as compared to High_notCNE bases, we examined the absolute SFS normalized by the number of base pairs and an estimated average mutation rate^30^ (**Methods**). We found that there are fewer SNPs at the Low_CNE bases relative to High_notCNE bases across all bins of allele frequencies (**Fig. 4b, Supplementary Fig. 10b,d,f**, comparing orange and purple bars). These results further suggest that the Low_CNE bases have experienced stronger purifying selection than the High_notCNE bases. We verified that similar results were obtained when controlling for differences in estimated background selection in different sets^31^ (**Methods, Supplementary Fig. 11**).

**Figure 4:**
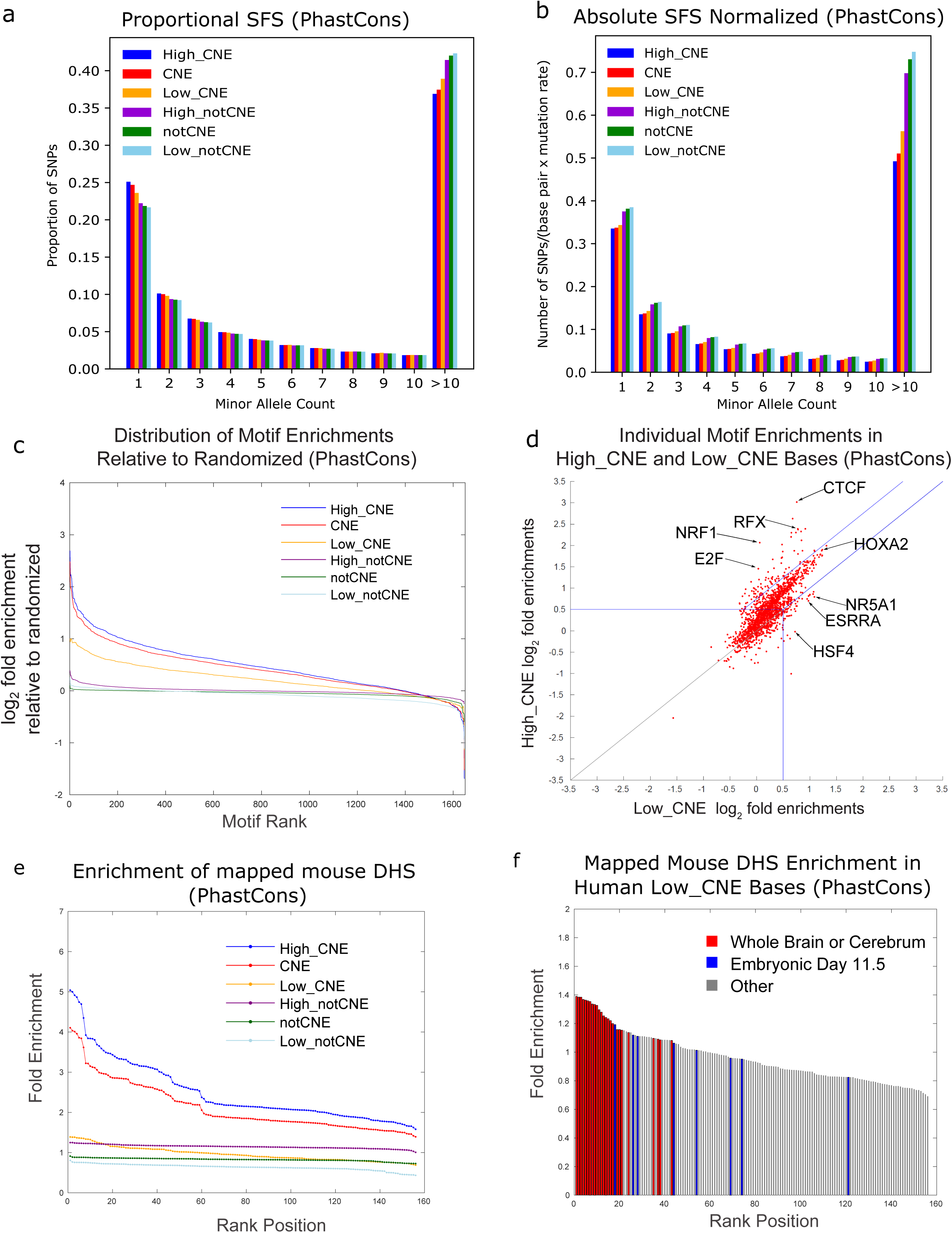
CNEP score’s relationship to human variation, TF sequence motifs, and DNase I Hypersensitive Sites in mouse. **(a)** The plot shows for PhastCons High_CNE, CNE, Low_CNE, High_notCNE, notCNE, and Low_CNE bases the proportional site frequency spectrum based on a set of 105 unrelated individuals in the YRI population in terms of # SNPs per base pair eligible for a SNP to be called, normalized for the number of sites with a variant in each set (**Methods**). The last column includes all SNPs with minor allele count greater than 10. **(b)** Similar plot to (a), except showing the absolute site frequency spectrum per base pair eligible for a SNP to be called normalized by estimated mutation rates. Corresponding plots for additional constrained elements can be found in **Supplementary Fig. 10** and plots controlling for difference in background selection can be found in **Supplementary Fig. 11**. Plots at higher thresholds of the CNEP score for notCNE bases can be found in **Supplementary Fig. 12. (c)** The plot shows the difference of the distribution of motif enrichments relative to the distribution for a randomized set of the motifs for the PhastCons High_CNE, CNE, Low_CNE, High_notCNE, notCNE, and Low_CNE bases. The x-axis is the rank position of the motif among the 1,646 motifs. The y-axis is the difference between the log_2_ fold enrichment based on the actual motif calls and the median log_2_ fold enrichment from three randomized versions at the same rank position (**Methods**). Similar plots for other constrained elements can be found in **Supplementary Fig. 13** and at other thresholds for defining notCNE high bases in **Supplementary Fig. 14. (d)** Scatter plot of individual motif enrichments. The x and y axes corresponds to the log_2_ fold enrichments in PhastCons Low_CNE and High_CNE bases respectively. The blue lines separate the three regions used for the GO enrichment analysis, High_CNE strongly preferred, High_CNE moderately preferred, and Low_CNE preferred, where at least one of the Low_CNE or the High_CNE log_2_ enrichment is greater than or equal to 0.5 (**Supplementary Table 6**). The gray line is the y=x line where both Low_CNE and High_CNE log_2_ enrichments are less than 0.5. Similar plots based on other thresholds of the CNEP score can be found in **Supplementary Fig. 15**. Selected motifs are labeled. **(e)** The distribution of enrichments for DNase I Hypersensitive Sites (DHS) from 156 experiments in mouse, where the sites are mapped to human and enrichments are computed relative to enrichments for a randomized DHS, for PhastCons High_CNE, CNE, Low_CNE, High_notCNE, notCNE, and Low_CNE bases (**Methods**). Similar plots for other constrained elements can be found in **Supplementary Fig. 16. (f)** A bar graph corresponding to the enrichments shown in (e) for Low_CNE bases. Bars are colored to indicate if the experiment is of whole brain or cerebrum (red), embryonic day 11.5 (dark blue), or neither (gray). Similar plots for other constrained elements can be found in **Supplementary Fig. 17**. A table of the enrichment values can be found in **Supplementary Table 7**.

We also observed that High_CNE relative to Low_CNE bases and High_notCNE relative to Low_notCNE bases had reduced common variation (**Fig. 4a,b, Supplementary Fig. 10**). However, these differences were generally smaller than the differences of CNE to notCNE bases. Additionally, we compared SFS of Low_CNE bases to subsets of High_notCNE bases that satisfied more stringent thresholds on the CNEP score (**Supplementary Fig. 12**), which suggested that the Low_CNE are under stronger purifying selection in humans than notCNE bases that receive substantially higher CNEP scores.

### Low_CNE bases show enrichments for TF binding motifs

Since human population genetics data still supported the importance of Low_CNE bases despite the low CNEP score, we investigated whether regulatory sequence motif analysis, which is cell type and condition invariant, provides evidence of a regulatory role for Low_CNE bases. For each constrained element set, we computed individual motif enrichments of Low_CNE bases relative to control motif instances from a compendium of 1,646 regulatory motifs, and did the same for the other five sets above (**Fig. 4c, Supplementary Fig. 13, Methods**). We then analyzed the distributions of these motif enrichments relative to those obtained from randomizing the motif instances. Both the Low_CNE and High_notCNE sets have motif enrichments that are above background, though less than High_CNE bases. The motif enrichments for the Low_CNE bases were substantially stronger than compared to the High_notCNE and notCNE bases. To place the motif enrichments of Low_CNE bases in additional context, we compared them to High_notCNE defined at more stringent thresholds of the CNEP score (**Supplementary Fig. 14**). Similar to what we saw with the SFS analysis, the Low_CNE bases had greater enrichment for motifs than High_notCNE at more stringent thresholds, with the specific thresholds depending on the constrained element set being considered.

To understand the specific motifs driving the overall motif distribution enrichments of the Low_CNE and High_CNE bases, we investigated directly the enrichments of individual motifs (**Fig. 4d, Supplementary Fig. 15, Supplementary Table 5**). While globally the High_CNE bases had stronger motif enrichments than Low_CNE bases, we did observe some motifs that showed enrichment for Low_CNE bases and for which the enrichment was greater than for High_CNE bases. We conducted a Gene Ontology (GO) enrichment for TFs corresponding to the set of motifs that had at least a log_2_ fold enrichment of 0.5 in High_CNE or Low_CNE bases broken down into three sets: ‘High_CNE strongly preferred’, ‘High_CNE moderately preferred’, and ‘Low_CNE preferred’ (**Fig. 4d, Supplementary Table 6, Methods**). TFs associated with ‘High_CNE strongly preferred’ motifs showed significant enrichment for protein dimerization activity and core-promoter GO terms. TFs associated with ‘High_CNE moderately preferred’ motifs showed enrichment for development related GO terms. Finally, TFs associated with ‘Low_CNE preferred’ motifs showed enrichment for lipid binding, signaling, and response to stimulus related GO-terms (corrected p-values <0.05). These results suggest that some of the CNE bases that are more difficult to predict from the epigenomic and TF binding data considered are associated with sites that might only be active in specific developmental stages or under specific stimuli.

### Low_CNE bases show some enrichments for mapped mouse DNase I hypersensitive sites

As mouse experiments can potentially have coverage of cell or tissue types and developmental stages not represented in human data, we investigated the extent to which DNase I hypersensitive sites (DHS) in mouse mapped to the human genome enriched for Low_CNE bases. Specifically, we analyzed a set of 156 DNase I hypersensitivity experiments from the mouse ENCODE project^32,33^. For each experiment, we mapped the set of mouse DHS to the human genome and did the same for datasets where we first randomized the location of the peaks in mouse (**Methods**). For each constrained element set, we then computed the enrichment of Low_CNE bases for the actual dataset relative to the randomized dataset, and also did the same for the other five sets above.

We found that for all element sets, the Low_CNE bases showed enrichment for the mapped DHS at least for a subset of the experiments (**Fig. 4e, Supplementary Fig. 16**). These enrichments were modest, not exceeding 2-fold for any DHS experiment or constrained element set, and were lower than for High_CNE bases. However, the enrichments were greater than for Low_notCNE and notCNE bases and comparable to High_notCNE bases for at least the most enriched experiments. We observed that the DHS experiments that tended to have the greatest enrichment for Low_CNE bases were for whole brain or cerebrum (**Fig. 4f, Supplementary Fig. 17, Supplementary Table 7**). For example, for PhastCons, the 21 DHS experiments with the greatest enrichment included 20 of the 25 experiments done on whole brain and cerebrum. The only other experiment in the top 21, was for the mesoderm conducted at day 11.5 in mouse embryos. These results provide evidence to suggest that a limited portion of Low_CNE bases are regulatory active in corresponding positions in available mouse samples, particularly related to the brain.

### Additional information to predict CNE bases in specialized human datasets

We conducted a retrospective analysis to gain insight into the extent to which additional data improves the predictive performance of CNEP and the nature of individual data sets that can offer additional marginal information predictive of CNE bases even after thousands of datasets are generated. Specifically, we generated another set of CNEP predictions based on a subset of 10,836 features that were available and accessible by 2015 (**Methods**). Using this subset of features we obtained lower AUC values for the ROC curve for each individual constrained element set with values in the range [0.75, 0.82] compared to [0.79,0.86] with the full set of features (**Supplementary Table 3**).

This suggests that some individual features may provide additional marginal information predictive of CNE bases even after considering more than ten thousand features. To identify such features, we defined the CNEP underestimation value for a set of peaks as the difference between the expected average CNEP score based on its overlap with constrained element calls and actual observed average value of the CNEP score for bases in the peaks (**Methods**). New datasets with a large amount of marginal additional information for CNE bases would have a large value for the CNEP underestimation and also cover many bases of the genome.

For most features the CNEP underestimate for a dataset was relatively small (<∼0.01) or only applied to relatively few bases, indicating that the additional marginal information in those datasets is limited (**Fig. 5, Supplementary Fig. 18-19, Supplementary Table 8**). Among the exceptions was a dataset of peak calls from a DNase I hypersensitivity experiment in spinal cord of a 59-day embryo that covered 71.6 million bases and had an underestimation value of 0.066. Other features based on DNase I hypersensitivity of embryonic spinal cord, brain, eye and retina also had notably high combinations of underestimate values and genome coverage. Additionally, some TF binding data sets in specialized cell types had underestimate values greater than observed in the embryonic DNase I hypersensitivity experiments, though covered a smaller fraction of the genome. We used the ChIP-atlas^26^ cell type class metadata annotations to determine if there were specific classes that had datasets significantly enriched with CNEP underestimate values greater than 0.02, restricted to those datasets covering at least 200kb (**Supplementary Table 9**). We found enrichment of datasets of pluripotent stem cell, pancreas, and neural to be the most significantly enriched (corrected p-value <0.01).

**Figure 5:**
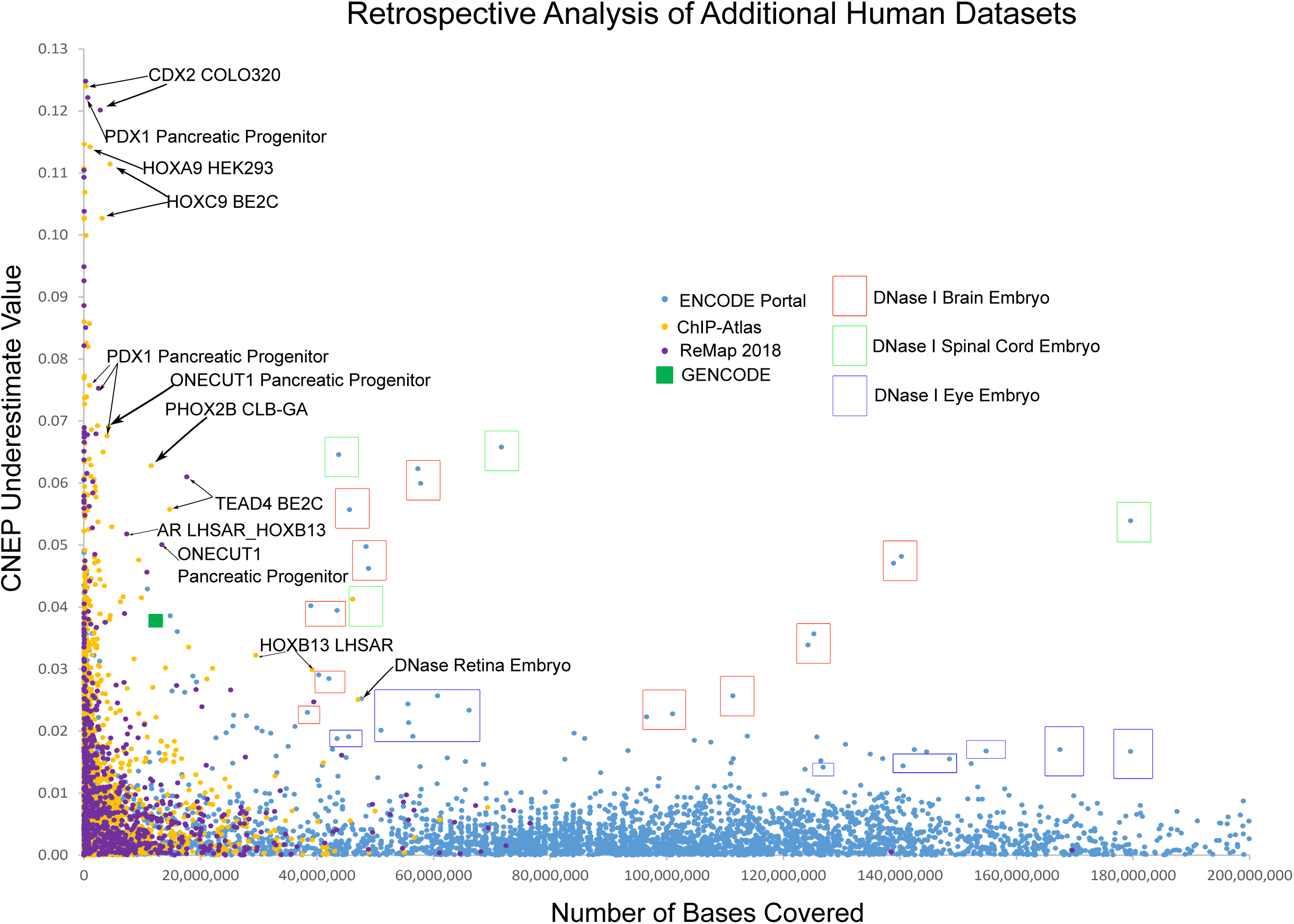
Retrospective analysis of information in additional human datasets. Scatter plot where each point corresponds to a dataset, the x-axis is the number of bases it covers, and the y-axis is the prediction underestimate value when using the CNEP score for prediction values. Selected datasets with a high combination of base coverage and underestimate values are labeled or placed in a box if they correspond to a DNase I hypersensitive experiment of embryonic brain, spinal cord, or eye. The color of the box corresponds to brain, spinal cord, or eye as indicated in the legend. The color and shape of the points are based on whether the point corresponds to a ChIP-atlas, ENCODE portal, or ReMap 2018 dataset or the set of new GENCODE exons between v19 and v28. Only datasets with a positive underestimate value are shown. Datasets covering more than 200 million base pairs are not shown, but all had an underestimate value of less than 0.01. Three datasets that had an underestimate value greater than 0.13 are not shown, but all covered less than 9,000 base pairs. Versions of these plots based on the input features to CNEP and based on shuffled versions of the additional datasets can be found in **Supplementary Fig. 19**.

We also compared the underestimate values and coverage of these datasets to new exons added to GENCODE between v19 and v28. New exons in GENCODE covered 12.3 million bases with an underestimate of 0.038. These values, while greater than those seen for many experiments, were still less than a single set of peak calls for both the TEAD4 and ONECUT1 TFs in terms of both genomic coverage and underestimate value and substantially less than some of the embryonic DNase I hypersensitivity experiments.

## Discussion

In this work, we developed and applied the Constrained Non-Exonic Predictor (CNEP) to provide a score for each base of the human genome that reflects the probability that the base overlaps a CNE from information in large-scale collections of epigenomic and TF binding data. We used information from more than 60,000 features derived from epigenomic and TF binding data spanning a wide range of cell and tissue types. We showed that CNEP outperformed baseline and related existing scores at predicting constrained non-exonic bases using epigenomic and TF binding data, but that there was still a substantial portion of CNE bases that were not well predicted. For example, 35% of CNE bases as defined by PhastCons had a CNEP score below the genome-wide average, while 23% of notCNE bases had a score above that average.

The CNEP score is designed to allow a researcher interested in gaining additional insights into a variant prioritized based on being in a constrained base to determine whether there are annotations within a compendium of epigenomic and TF binding data to explain the constraint. By defining the CNEP score without conservation annotations as features, we were also able to derive the complementary CSS-CNEP. The CSS-CNEP uses the combination of the ConsHMM conservation state and constrained element annotations to predict the CNEP score at a base. This allows one to differentiate among the CNE bases, which are more likely lacking predictive epigenomic and TF binding annotations due to false calls of constraint from those for which informative experiments for the base have not been conducted.

To better understand the nature of CNE bases that received a low CNEP, as well as notCNE bases that received a high CNEP score, we conducted additional analyses. Using human population genetic variation data, we provided evidence to suggest that the Low_CNE bases are under constraint in humans, though to a lesser extent than the High_CNE bases. Consistent with the trends of the human population variation data, regulatory sequence motifs also showed enrichment in Low_CNE bases, though to a lesser extent than in the High_CNE bases. High_notCNE bases showed greater enrichment for regulatory motifs and less genetic variation compared to Low_notCNE bases, though less than Low_CNE bases. A subset of High_notCNE bases might correspond to bases that are under evolutionary constraint in humans, but not actually in a constrained element call, which was supported by the conservation state enrichments for High_notCNE bases. Another subset of High_notCNE bases may correspond to adaptive and recently evolved bases with a potentially important regulatory role that share epigenomic marks and TF binding patterns associated with CNE bases.

Through a retrospective analysis, we showed additional data led to modest overall improvements in the CNEP predictions. Datasets with substantial additional information were limited and we saw an enrichment of datasets with relatively informative additional marginal information categorized as being from pluripotent stem cell, pancreas, or neural cell type classes. It is possible different types of assays or cell types and conditions distinct from those considered here would enable improved prediction of CNE bases.

While the CNEP score was a relatively effective predictor of CNE bases from epigenomic and TF binding compared to other approaches, a portion of CNE bases remains difficult to predict from such compendiums. The functional importance of a subset of such bases is supported by orthogonal evidence, thus also highlighting the remaining challenge to a comprehensive understanding of the non-exonic genome.

## Methods

### Availability of CNEP and CSS-CNEP scores and CNEP software

The CNEP and the CSS-CNEP scores and software are available from https://github.com/ernstlab/CNEP.

### Genome assembly and gene annotations

All predictions and analysis were done on human genome assembly hg19 and were restricted to chr1-22 and chrX. For gene annotations we used the GENCODE v19 annotations obtained from ftp://ftp.sanger.ac.uk/pub/gencode/Gencode_human/release_19/gencode.v19.annotation.gtf.gz. Exon annotations include exon bases that are non-coding.

### Constrained element sets

We used four different constrained element sets based on the PhastCons^5^, GERP++^2^, SiPhy-omega, and SiPhy-pi^3,4^ methods. The PhastCons constrained elements were based on the human hg19 100-way vertebrate alignment and obtained from the UCSC genome browser^29^. The SiPhy-omega and SiPhy-pi elements were called based on a 29-way mammalian alignment and were the hg19 version obtained from https://www.broadinstitute.org/mammals-models/29-mammals-project-supplementary-info. The GERP++ elements were called based on the mammalian subset of the UCSC genome browser hg19 46-way vertebrate alignment obtained from http://mendel.stanford.edu/SidowLab/downloads/gerp/.

### Epigenomics and TF binding features

We used 63,741 binary features defined from functional genomics data. The sources of the features are found in **Supplementary Table 1** and a list of features can be found in **Supplementary Table 2**. The features were derived from ChIP-seq data of histone modifications, TFs including general factors, DNase I hypersensitivity data, and FAIRE data. The data were from the ENCODE consortium^11,34^, Roadmap Epigenomics consortium^14^, the ReMap public dataset^25,27^, or the ChIP-atlas^26^.

In total 58,484 features were based on peak calls. For these features, the data was encoded as a ‘1’ if the corresponding base overlapped a peak and ‘0’ otherwise. The peak calls for the Roadmap Epigenomics data was based on the unconsolidated datasets. For peak calls from the ENCODE consortium, we combined sets of peak calls from the second phase of the ENCODE project^11^ with peak calls available from the ENCODE portal^34^. For the ENCODE portal data we downloaded all files for the ChIP-seq or DNase-seq assay available in narrowPeak or broadPeak format for hg19 on May 11, 2018 produced by the ENCODE project from https://www.encodeproject.org/. For the ReMap database, which is a reprocessing of ChIP-seq data of TFs from the Gene Expression Omnibus and ArrayExpress, we used the peaks in the hg19 files restricted to the ‘Public’ data (non-ENCODE) and had features based on both the 2015 and 2018 versions of the database. For the ChIP-atlas, which is a reprocessing of Sequence Read Archive data, we used all peaks called at the 10^−5^ threshold for hg19 available from http://dbarchive.biosciencedbc.jp/kyushu-u/hg19/eachData/bed05/ on May 11, 2018. We note that some of the ChIP-atlas datasets were generated by the ENCODE or Roadmap Epigenomics project, but processed differently.

We also had 5,215 features defined based on chromatin state calls from three different ChromHMM models^35^. The three models were: (1) a 15-state model defined across 9-ENCODE cell types based on eight histone modifications and CTCF; (2) the 15-state ‘core’ model based on 5-histone modifications defined across 127-reference epigenomes based on consolidated data processed by the Roadmap Epigenomics consortium (111 reference epigenomes were derived from data produced by the Roadmap Epigenomics project and 16 from the ENCODE project); (3) a 25-state model based on imputed data for 12-chromatin marks (10 histone modifications, H2A.Z, and DNase I hypersensitivity) defined across the same 127 reference epigenomes^36^. For each model, we had a separate feature for each chromatin state and cell type or reference epigenome combination. A feature value was encoded as a ‘1’ if a base overlapped the chromatin state in the cell type or reference epigenome and a ‘0’ otherwise.

Additionally, we had 42 features defined based on the position of Digital Genomics Footprints^14,37^. For these features the data was encoded as a ‘1’ for those bases overlapping a footprint and a ‘0’ otherwise.

### CNEP method

The CNEP scores are generated by first training an ensemble of logistic regression classifiers. For a given constrained element set and a set of binary functional genomics features, CNEP trains logistic regression classifiers to discriminate between bases in a constrained element that are outside of all exons as a positive set from all other bases, including exons, as a negative set. For generating CNEP scores on one chromosome based on one constrained element set, CNEP trained ten logistic regression classifiers using different 1,000,000 randomly sampled positions from the other 22 chromosomes. We repeated this for each of the constrained element sets and 23 chromosomes thus training in total 920 logistic regression classifiers in parallel. CNEP used the Liblinear v.2.1^38^ software to train the logistic regression classifiers using L_1_ regularization (-s 6) with a bias term (-B 1), with the default regularization parameter value of 1 (-c 1). The exclusive use of binary features allowed us to make efficient use of the sparse representation of the data in Liblinear. For generating genome-wide predictions based on one constrained element for each chromosome, CNEP computed and averaged the probabilistic predictions from its ten corresponding logistic regression classifiers and then outputted the predictions to the nearest 0.001 value. To generate the CNEP score we then averaged the outputted predictions based on each of the four constrained element sets.

### Computing actual observed and expected average CNEP scores for features and genome-wide

For computing the actual observed average CNEP score for a feature, we computed the average CNEP score in all bases in the genome where the feature was defined as being present. For computing the expected average CNEP score for a feature, we computed the average over the four constrained element sets of the number of bases for which the feature was present and overlapped a constrained non-exonic element divided by the total number of bases in which the feature was present. We computed the genome-wide observed and expected CNEP scores the same way except all bases in the genome were included.

### Analysis of CNEP score’s relationship to Roadmap Epigenomics DNase I hypersensitive peak coverage

For computing the relationship between CNEP score and average fraction of Roadmap Epigenomics DNase I hypersensitivity experiments in a peak (**Fig. 2c**), we used 350 narrowPeak call files with ‘ChromatinAccessibility’ in the file name available from http://egg2.wustl.edu/roadmap/data/byFileType/peaks/unconsolidated/narrowPeak/. For each value of the CNEP score computed to the nearest 0.001 and covering at least 1000 bases, we took all bases in the genome having that score and determined the average fraction of the 350 experiments in which the bases are overlapped by a peak call.

### Baseline and related score comparisons

For comparing the predictive performance of the CNEP score, the baseline score based on Roadmap Epigenomics DNase I hypersensitivity data was computed by computing the number of times a base was overlapped for the set of data above. For the all DNase I baseline evaluation, we expanded the counts to be based on a set of 4,522 features annotated to be peaks from DNase I hypersensitive experiments. We also generated a baseline based on the number of times any of the 63,741 features overlap a base. We also compared to each of those features individually. In addition we compared to two other existing related scores, the ‘Position-wise aggregated conservation-associated activity scores’^23^, that we obtained from: https://noble.gs.washington.edu/proj/encyclopedia/caas.bed.gz and the FitCons2 cell-type integrated score^21^ that we obtained from http://compgen.cshl.edu/fitCons2/hg19/H1/E999-sco.bw

### Analysis of CNEP score’s relationship to chromatin states

For the analysis of the relationship between CNEP score and chromatin state annotation (**Fig. 2d**), we used the 25-state ChromHMM chromatin state annotations defined across 127 epigenomes based on imputed data for 12-chromatin marks^36^. For each CNEP score, which were computed to the nearest 0.001 value, we took all bases in the genome having that score and determined the average fraction of the 127 epigenomes in which each of the 25-states overlapped the base. We then stacked bar graphs with fractions starting from the state with the greatest state number (25_Quies) to the lowest state number (1_TssA) with the state numbers and colors from Ref. 36. In the plot, we did not differentiate between different states that were previously given the same color and thus the graph provides information on the 14-state groups that were colored differently.

### Defining sets of bases for analyses

For each of the four constrained element sets considered, we defined the following set of bases used in some analyses: (1) CNE – bases in a constrained element that do not overlap a GENCODE exon; (2) Low_CNE – bases in a constrained element that have a CNEP score less than or equal to the CNEP mean, 0.0419; (3) High_CNE – bases in a constrained element that have a CNEP score greater than 0.0419; (4) notCNE – bases not in a constrained element and do not overlap a GENCODE exon; (5) Low_notCNE – bases not in a constrained element and do not overlap a GENCODE exon and have a CNEP score less than or equal to 0.0419; (6) High_notCNE – bases not in a constrained element and do not overlap a GENCODE exon and have a CNEP score greater than 0.0419 (**Supplementary Table 3**).

### Conservation state analysis

The ConsHMM conservation state annotations were the 100-conservation state annotations for hg19 from Ref. 24. These conservation state annotations were defined on the same 100-way vertebrate alignment for which the PhastCons bases we used were defined. Fold enrichments for CNE, Low_CNE, notCNE, and High_notCNE bases in the conservation states were computed using the OverlapEnrichment command of ChromHMM v1.17 with the options ‘-b 1 -lowmem’ specified. Conservation state assignments to chrY were excluded from the background in this analysis, as the CNEP scores were not defined on this chromosome. The per state ROC and AUC values for the CNEP score were computed by considering a positive base a CNE in a specific conservation state and a negative base any base in the genome that was not in a CNE. Bases in the CNE set that were in a different conservation state were excluded when generating the ROC and computing the AUC. The ROC and AUC based on extending exons 200bp in each direction was computed in the same way, except first adjusting the exon start and end positions.

### Conservation Signature Score by CNEP (CSS-CNEP)

To compute the Conservation Signature Score by CNEP (CSS-CNEP), for each chromosome we first computed the average CNEP score for each combination of the 100 conservation states and four constrained elements within non-exonic bases on all other chromosomes. We then for each base on the target chromosome used the average corresponding to the combination of conservation state and constrained element sets present at the base. In total, there was 1600 such combinations.

### Evaluation of predictions of Low_CNE bases among CNE bases with comparative genomics features

We evaluated the ability of CSS-CNEP and other comparative genomic scores and annotations to predict among CNE bases those that will receive a low CNEP score. Specifically for this evaluation, we split the CNE bases into two sets, the Low_CNE and High_CNE bases, and treated the Low_CNE bases as positives and High_CNE bases as negatives in the evaluation. We followed a previous evaluation approach where we randomly split 200kb genome-segments into two halves, one used for training and the other used for testing^24^. The precision-recall for ConsHMM states was based on decreasing order of ConsHMM states enrichment for Low_CNE bases with a background of all CNE bases on the training data, and then using that order to evaluate the cumulative precision-recall on the test data. We formed predictions for the CSS-CNEP, PhyloP^39^, GERP++^2^, and PhastCons^5^ scores by ordering bases from lowest to highest value according to the score. For the PhyloP, GERP++, and PhastCons scores we also followed the procedure of Ref. 24 and formed 400 equal sized bins of the score and then repeated the procedure used for the ConsHMM conservation states treating a bin as if it was a state. For each constrained element set other than the one used to define the CNE bases in the evaluation, we computed the precision and recall for a prediction based on bases not overlapping the constrained element set.

### Positional enrichment analysis relative to exons and CNE bases

For computing the enrichment of Low_CNE bases in proximity to exons we computed for each base in the genome the distance to the nearest base of any exon. The enrichment of Low_CNE at a specific distance to the nearest exon was defined as the ratio of the fraction of Low_CNE bases whose nearest exon was at that distance to the fraction of all bases in the genome whose nearest exon was at that distance. A similar set of enrichments was computed for High_notCNE bases in relation to their distance to the nearest CNE base.

### Human variation analysis

The human variation analysis was conducted on a set of 105 unrelated individuals from the Yoruba in Ibadan (YRI) population part of the 1000 Genomes Project. We focused on this population for analyzing the effects of selection since it is associated with greater genetic diversity and has a simpler demographic history than non-African populations^40^. We selected high-quality sites by applying a mask from 1000G where a site was defined as high quality if its depth (DP) is within 1.5x the mean DP across all sites^41^. For this analysis, we restricted it to the autosomes and variant calls that were bi-allelic. For each set of coordinates analyzed, we computed a count c_*n*_ of how many of the variants occurred in exactly *n* individuals for each value of *n*=1,…,10 (low and intermediate frequency variants) and also a count c_*>10*_ of how many occurred in greater than 10 individuals (common variants). We then computed the proportional site frequency spectrum (SFS) as each of these individual counts divided by the sum of all of the counts. We assessed the statistical significance between pairs of coordinate sets by applying a chi-square test to the 11 count values.

The absolute SFS contains the numbers (rather than proportions) of SNPs at particular minor allele counts. Because the count of SNPs is affected by the number of base pairs analyzed (more base pairs would lead to more SNPs) as well the mutation rate (higher mutation rates lead to more SNPs), we normalized for both of these factors. To do this, first we obtained mutation rate estimates from http://mutation.sph.umich.edu/hg19/^30^. We associated each base with a single mutation rate by averaging its three mutation rates, each corresponding to a mutation from the reference nucleotide to an alternative nucleotide. For a coordinate set, we computed the sum of the mutation rates at all bases that were high quality sites as defined above and had a mutation rate available. This sum is equivalent to the number of base pairs analyzed in a coordinate set times their average mutation rate. We computed the unnormalized count values as described above for the proportional SFS except excluded positions that did not have mutation rates available. We then divided these counts by the sum of the mutation rates.

To compute SFS controlled for background selection we used the version of *B*-values in hg19 as part of the CADD annotation set, which are based on the *B*-values from Ref. 31. For the proportional SFS, we reweighted variant calls in each coordinate set so that the *B*-value distribution was effectively the same as the distribution of *B*-values at all non-exonic bases with a variant call. This analysis was restricted to non-exonic variants that had an estimated *B*-value available. The weighting for a variant with a *B*-value, *x*, was p_a_(*x*)/p_s_(*x*) where p_a_(*x*) and p_s_(*x*) are the proportions of variants with the *B*-value *x* among all variants considered and the subset in the coordinate set, respectively. For the absolute SFS density normalized by its average mutation rate, we reweighted bases in each coordinate set so the *B*-value distribution was effectively the same as the distribution of *B*-values at all non-exonic bases. This analysis was restricted to non-exonic bases that were in a high quality site and had both estimated mutation rates and *B*-values available. The weighting for a base with a *B*-value, *x*, was p_c_(*x*)/p_s_(*x*) where p_c_(*x*) and p_s_(x) are the proportions of variants with the *B*-value *x* among all bases considered and the subset in the coordinate set respectively. The weighting was used in both counting variants and the sum of the mutation rates.

### Motif enrichment analysis

For the motif enrichment analysis (**Fig. 4c,d, Supplementary Fig. 13-15, Supplementary Table 5**), we used motif instances from http://compbio.mit.edu/encode-motifs/matches-with-controls.txt.gz^42^. We used motif instances for a set of 1,646 motifs that excluded motifs that were in the compendium based on being discovered from ENCODE ChIP-seq data, so that the set of motifs we analyzed were independent of the features provided to CNEP. The motif instances were called outside of coding, 3’ UTR, and repetitive regions and called independent of conservation. For each motif, there were also a set of corresponding control motif instances called^42^, which control for biases from sequence composition or background. To compute the enrichment of a specific motif in a target set of bases, we computed the ratio of the fraction of motif instance bases that also overlapped the target set to the fraction of corresponding control motif instance bases that also overlapped the target set. For each of the four constrained element sets, these enrichments for individual motifs were reported for High_CNE and Low_CNE bases (**Fig. 4d, Supplementary Fig. 15, Supplementary Table 5**). For the analyses of the distribution of motif enrichments for a target set, we generated three randomized versions of the motif instance calls with controls. To generate a randomized version for each chromosome we performed column-wise random permutations where one column is the motif identifier and the other column contains the motif coordinates. For each target set considered, we computed the distribution of motif enrichments on each of the randomized motif instances, using the same procedure as the actual motif instances. We then ordered the enrichments separately for the actual and three randomized datasets. At each ranked position in the ordering we took the difference between the log_2_ value of the actual enrichment and the log_2_ value of the median enrichment from the three randomized datasets.

### Motif set Gene Ontology analysis

We conducted Gene Ontology (GO) enrichment analysis for the TFs corresponding to three subsets of motifs that had at least a log_2_ fold enrichment of 0.5 in High_CNE or Low_CNE bases. The three subsets were: (i) ‘High_CNE strongly preferred’ - those motifs for which difference for High_CNE and Low_CNE bases was greater than 0.75; (ii) ‘High_CNE moderately preferred’ - the difference was between 0.75 and 0; (iii) ‘Low_CNE preferred’ - which had a greater enrichment for Low_CNE bases than High_CNE bases. GO enrichment was conducted using the STEM software v.1.3.11 with default settings (**Supplementary Table 6**)^43^. The Gene Ontology and human gene annotations were downloaded using the STEM software on September 17, 2017. We used as a base set all TFs corresponding to a motif in the compendium. The corresponding TF for a motif was taken to be the portion before the ‘_’ in the motif ID.

### Mouse DNase I hypersensitive site enrichment analysis

For the mouse DNase I hypersensitivity site (DHS) analysis (**Fig. 4e,f, Supplementary Fig. 16-17, Supplementary Table 7**), we used the 156 narrowPeak files from the University of Washington mouseENCODE group available from http://hgdownload.soe.ucsc.edu/goldenPath/mm9/encodeDCC/wgEncodeUwDnase/ and http://hgdownload.soe.ucsc.edu/goldenPath/mm9/encodeDCC/wgEncodeUwDgf/^32,33^. We also generated a randomized version of each set of DHS by randomly selecting a different position for each DHS in the original file on the same chromosome. We lifted over both the real and randomized versions of the DHS files from mm9 to hg19 using the liftOver tool from the UCSC genome browser with the options ‘-bedPlus=3 -minMatch=0.00000001’. The lower value for the minMatch parameter enables a more permissive mapping of peaks from mouse to human and thus an enrichment estimate that is more reflective of a background that includes all mouse DHS. For both the real and randomized version of each set of DHS, for each of the four constrained element sets considered, we computed enrichments for the six target sets: CNE, High_CNE, Low_CNE, notCNE, High_notCNE, and Low_notCNE. Enrichments for each set of DHS were computed by taking the ratio between the fraction of bases in human covered by a DHS that are in the target set to the fraction of bases in the genome that are in the target set. The reported enrichment for an experiment is the ratio of this enrichment for the real DHS compared to the corresponding enrichment for the randomized DHS.

### Retrospective analysis on information in additional human datasets

For the retrospective analysis of additional information in human datasets, we held out from training all features from ChIP-atlas^26^, ReMap 2018^27^, the ENCODE portal^11,34^. The version of the CNEP score that was generated with data available by 2015 was based on 10,836 features and included ENCODE consortium during its second phase^11^, Roadmap Epigenomics consortium^14^, or part of the ReMap 2015 public dataset^25^ (**Supplementary Table 1**). We did not exclude any dataset for being based on the same experiment as used to generate the CNEP predictions. We excluded files that did not have any peaks called on the chromosomes we considered.

For the analysis with updated GENCODE exon annotations we used release 28 mapped to hg19/GRCh37 available from ftp://ftp.ebi.ac.uk/pub/databases/gencode/Gencode_human/release_28/GRCh37_mapping/gencode.v28lift37.annotation.gtf We excluded any base in an exon from release 19. For generating the shuffled data we used the shuffleBed command of BEDTools^44^.

For computing the cell type class enrichments, we used STEM software v.1.3.11 with user provided annotations treating each dataset as if it was a gene and the cell type class as the annotation category (Ernst and Bar-Joseph, 2006). The foreground for the enrichment were those ChIP-atlas datasets with peaks covering at least 200kb and having an underestimate value greater than 0.02. The background set for the enrichment analysis was all ChIP-atlas datasets with peaks that covered at least 200kb. We used default settings except changed the minimum number of genes parameter to 1 and multiple hypothesis testing correction to ‘Bonferroni’.

## Acknowledgements

We thank Xiaorui Fan and Petko Fiziev for useful discussions and assistance. We acknowledge funding from US National Institutes of Health grants DP1DA044371, R01ES024995, U01HG007912 and U01MH105578 (J.E.), T32CA201160 (A.S.), and R35GM119856 (K.E.L.), US National Science Foundation CAREER Award #1254200 (J.E.), a Kure-IT award and an Alfred P. Sloan Fellowship (J.E.). We acknowledge the ENCODE and Roadmap Epigenomic consortia for generating some of the data used and making it available pre-publication.

