## Supplementary Figures for "Identification and characterization of constrained non-exonic bases lacking predictive epigenomic and transcription factor binding annotations"

| | CNEP (average) | CNEP SiPhy-PI only | CNEP SiPhy-omega only | CNEP PhastCons only | CNEP GERP++ only |
|---|---|---|---|---|---|
| CNEP (average) | | 0.96 | 0.98 | 0.96 | 0.98 |
| CNEP SiPhy-PI only | | | 0.93 | 0.88 | 0.91 |
| CNEP SiPhy-omega only | | | | 0.93 | 0.93 |
| CNEP PhastCons only | | | | | 0.92 |
| CNEP GERP++ only | | | | | |

Supplementary Figure 1

a

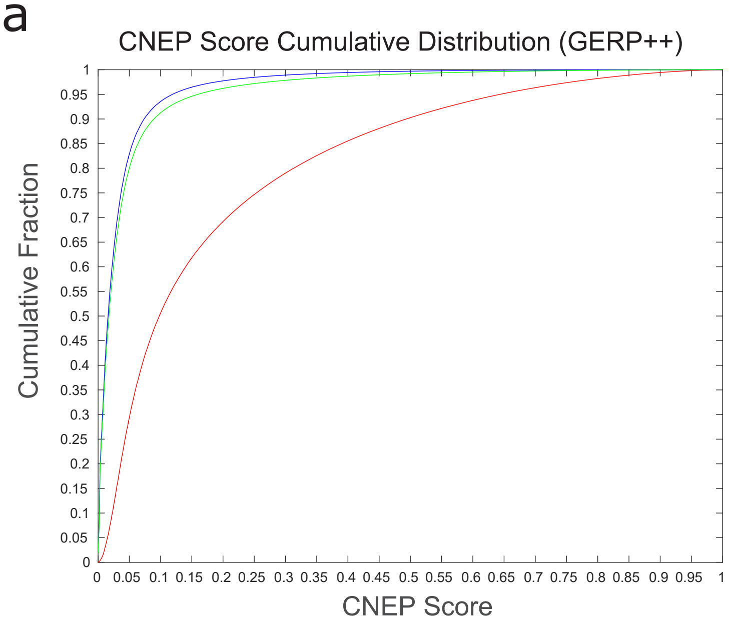


b

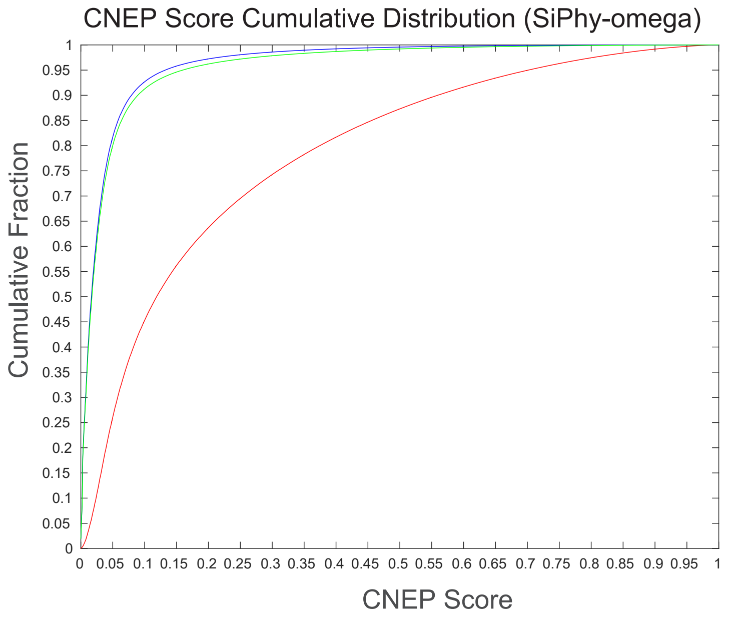


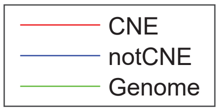


c

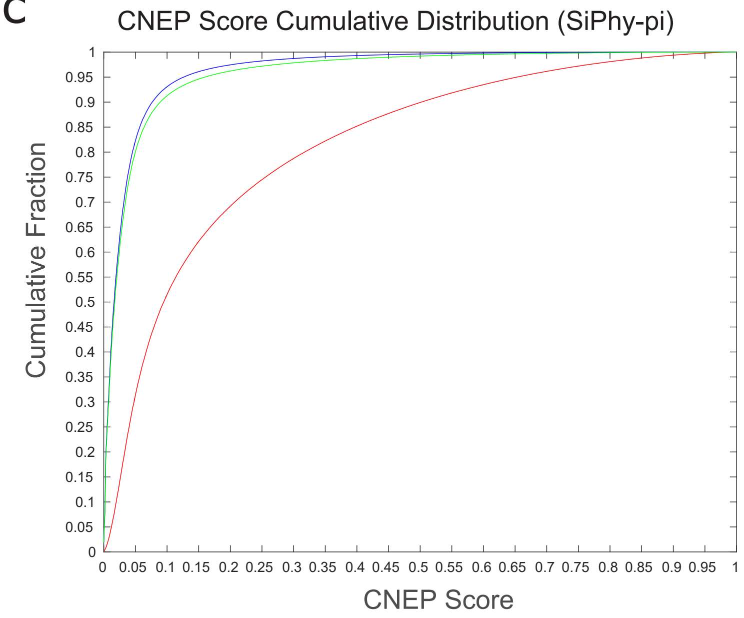


d

Genome Coverage %

| set | GERP++ | PhastCons | SiPhy-omega | SiPhy-pi |
|---|---|---|---|---|
| High_CNE | 4.1 | 2.5 | 2.4 | 3.2 |
| Low_CNE | 1.3 | 1.3 | 0.6 | 1.1 |
| CNE | 5.3 | 3.9 | 3.0 | 4.4 |
| High_notCNE | 19.6 | 21.1 | 21.3 | 20.4 |
| Low_notCNE | 71.1 | 71.0 | 71.7 | 71.2 |
| notCNE | 90.7 | 92.1 | 93.0 | 91.6 |

e

Non-exonic bases

| | CNEP below average (<= 0.0419) | CNEP above average (> 0.0419) |
|---|---|---|
| CNE | Low_CNE | High_CNE |
| notCNE | Low_notCNE | High_notCNE |

Supplementary Figure 2

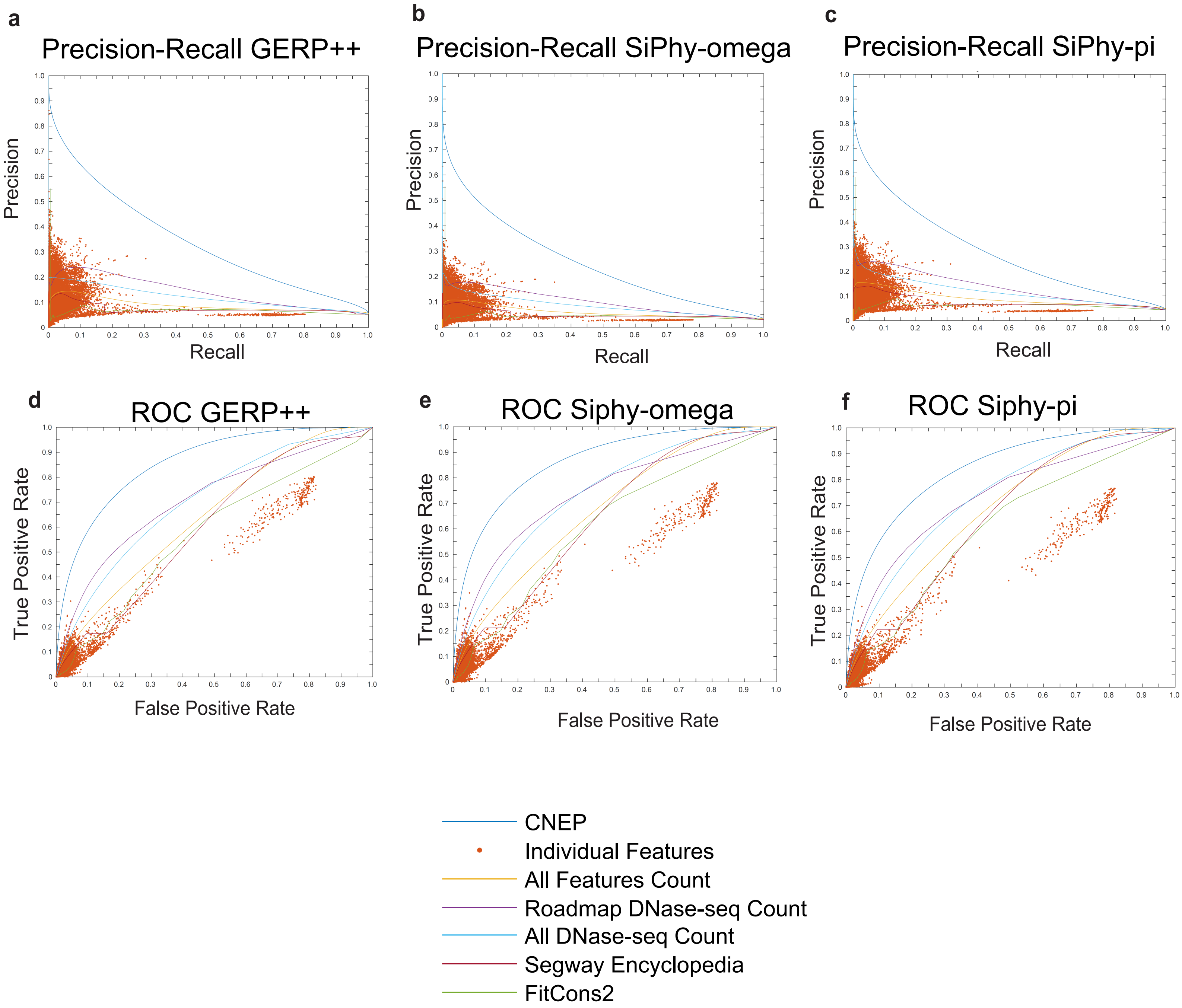


Supplementary Figure 3

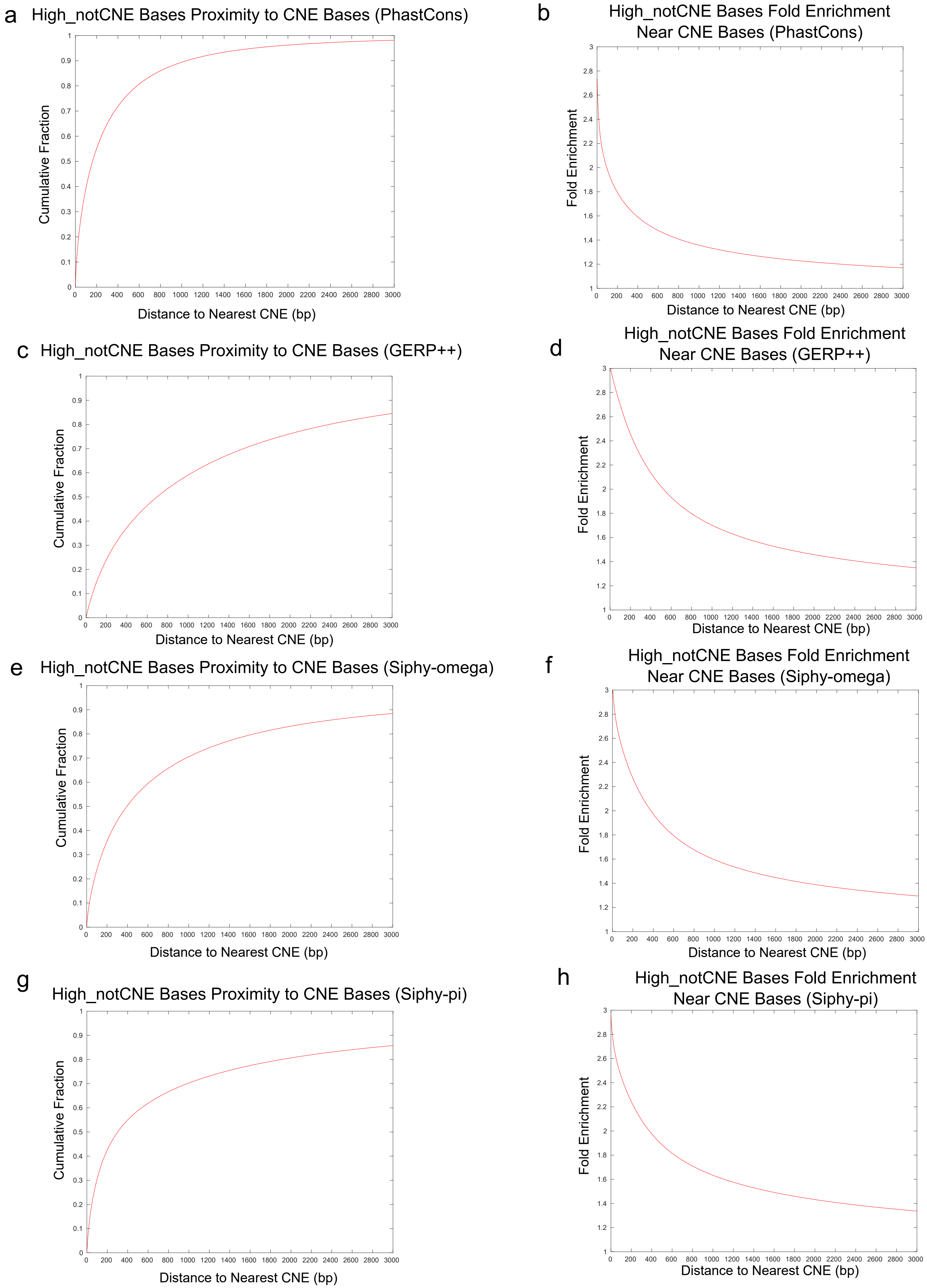


Supplementary Figure 4

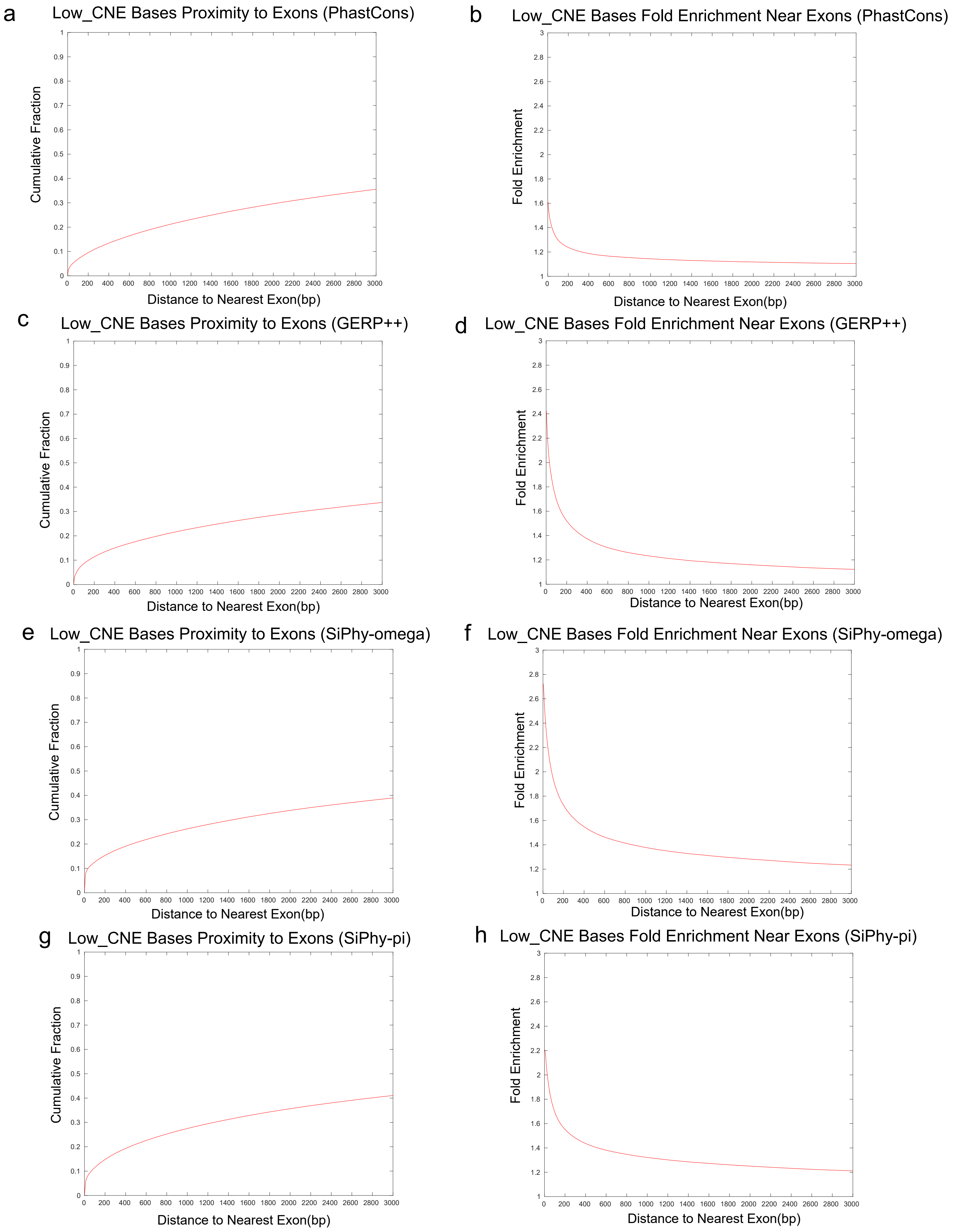


Supplementary Figure 5

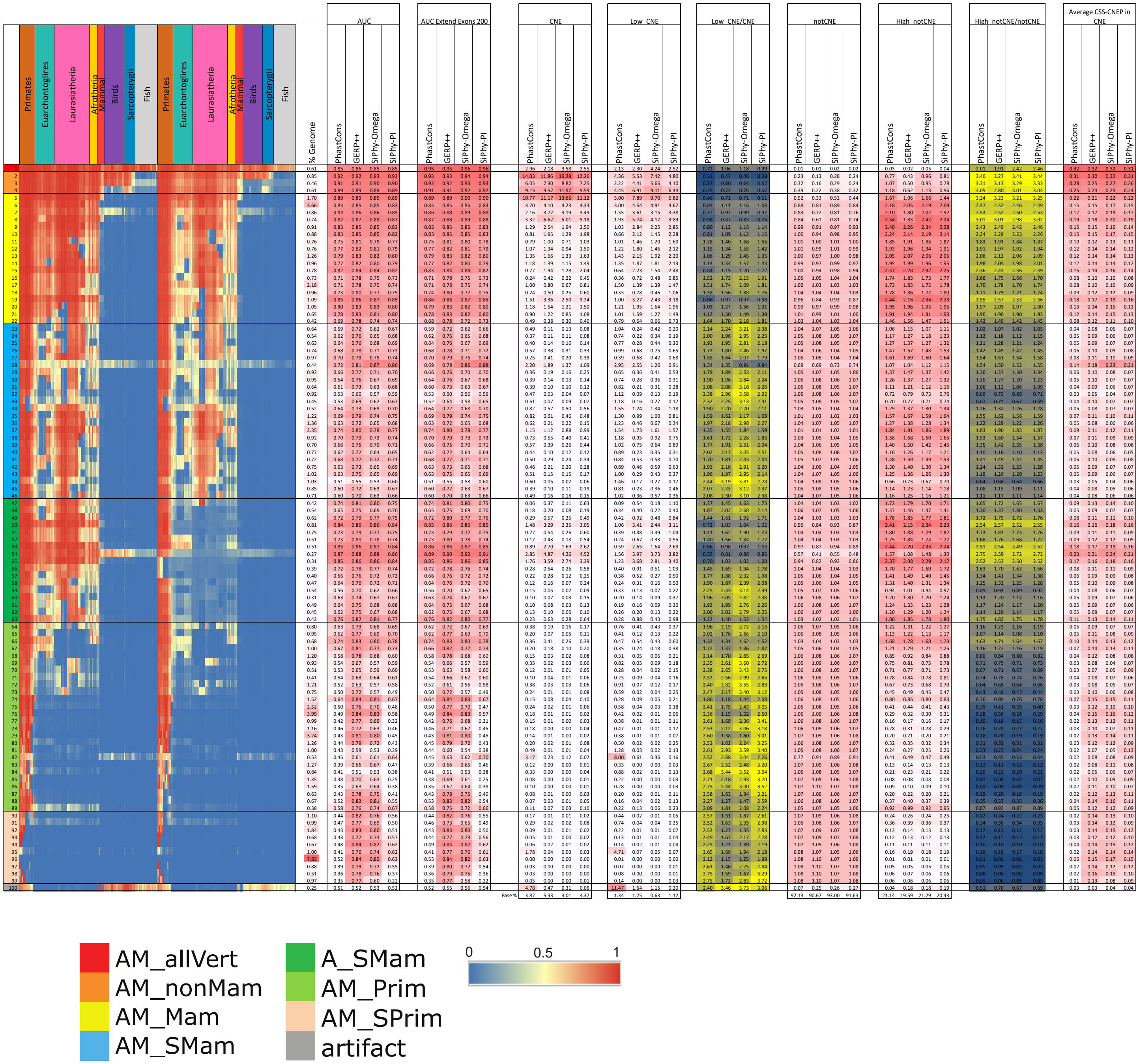


Supplementary Figure 6

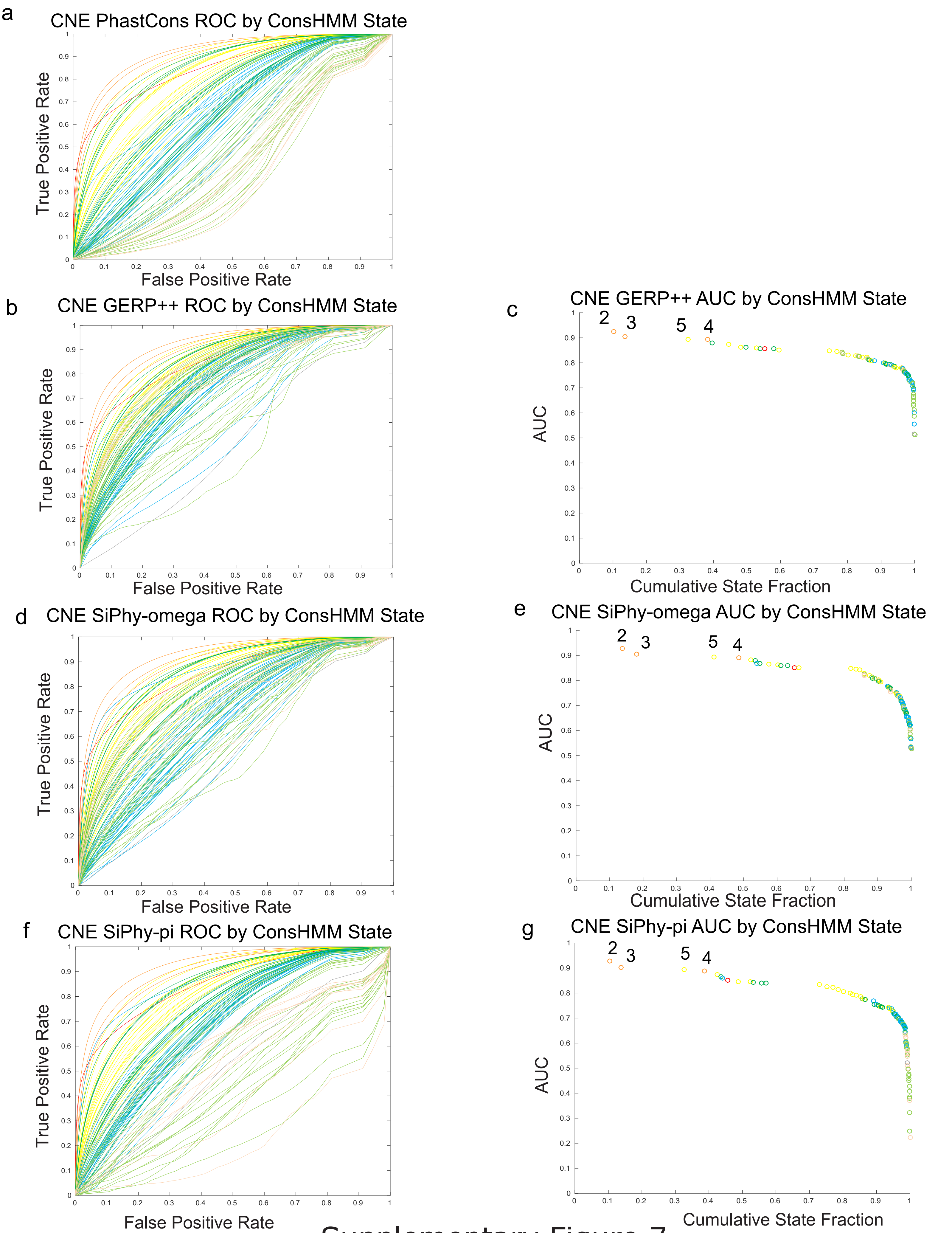


Supplementary Figure 7

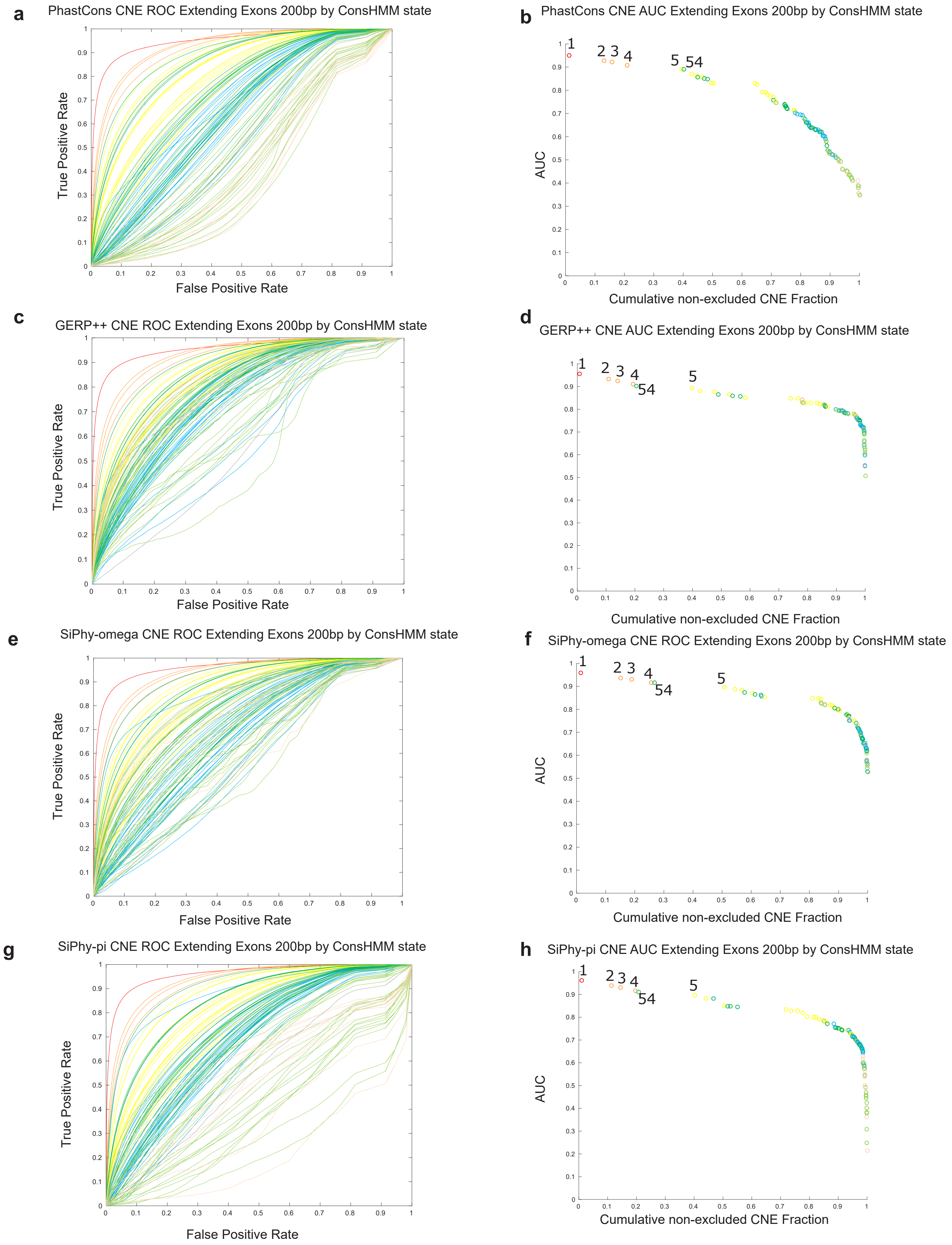


Supplementary Figure 8

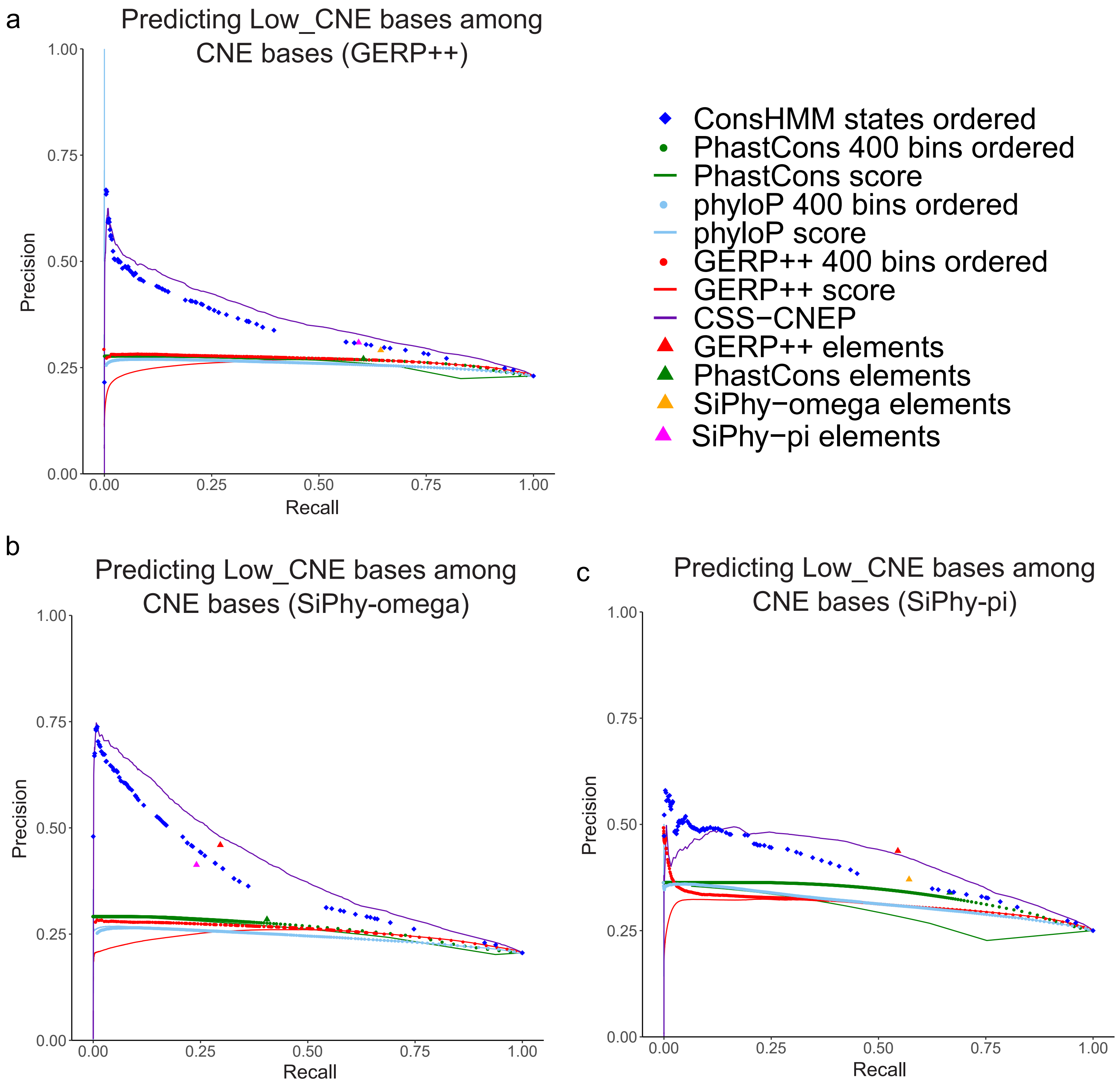


Supplementary Figure 9

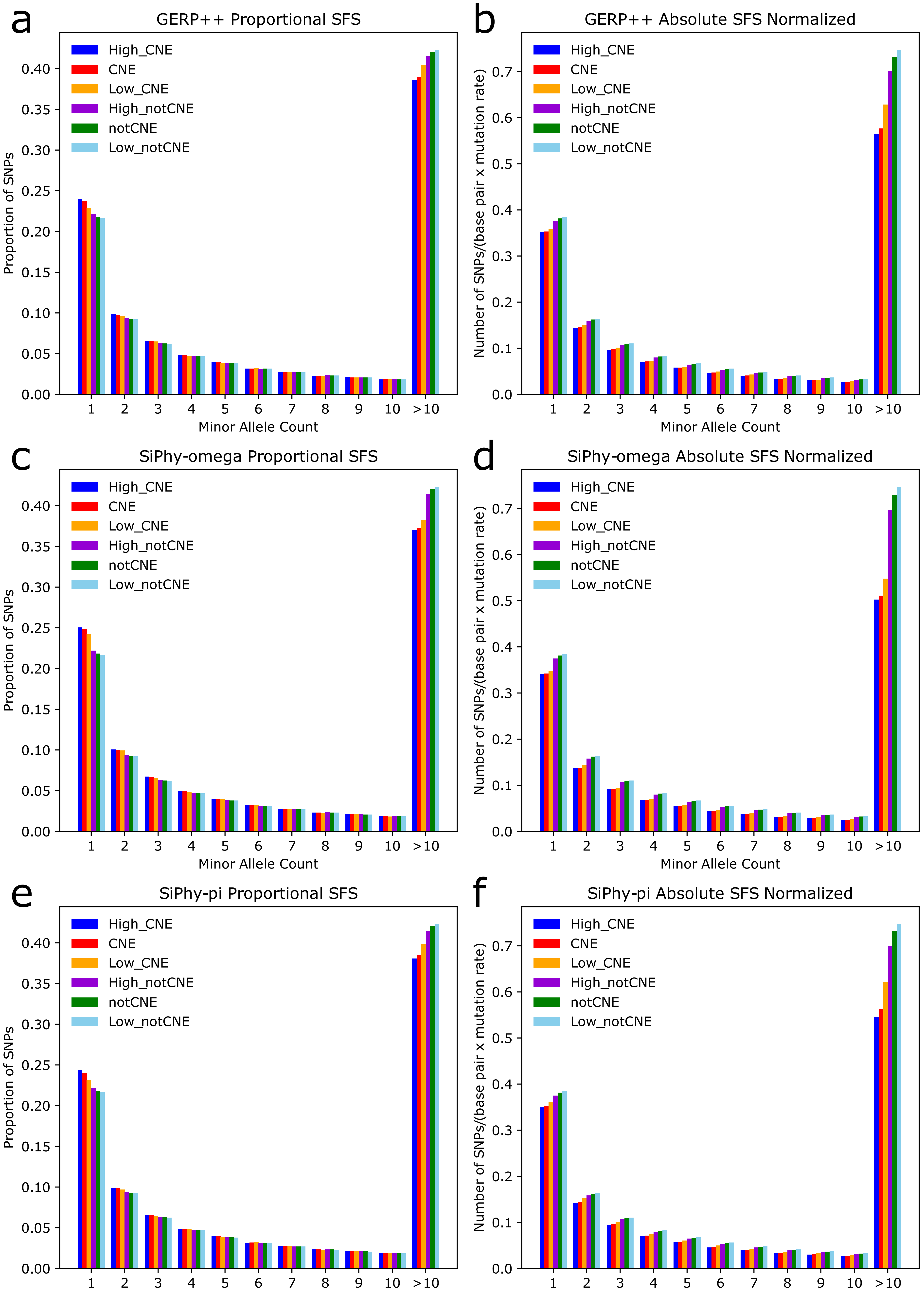


Supplementary Figure 10

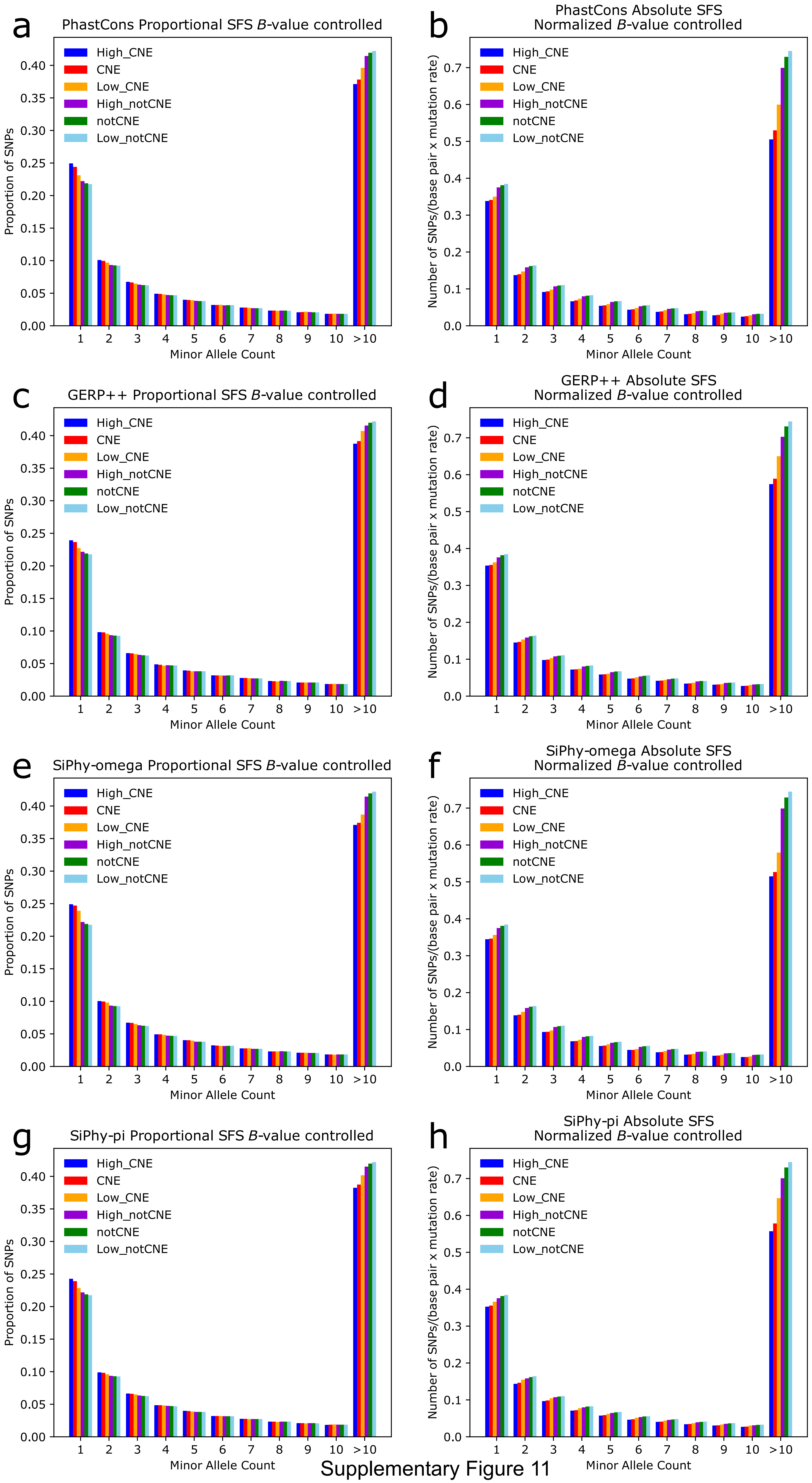

Supplementary Figure 11

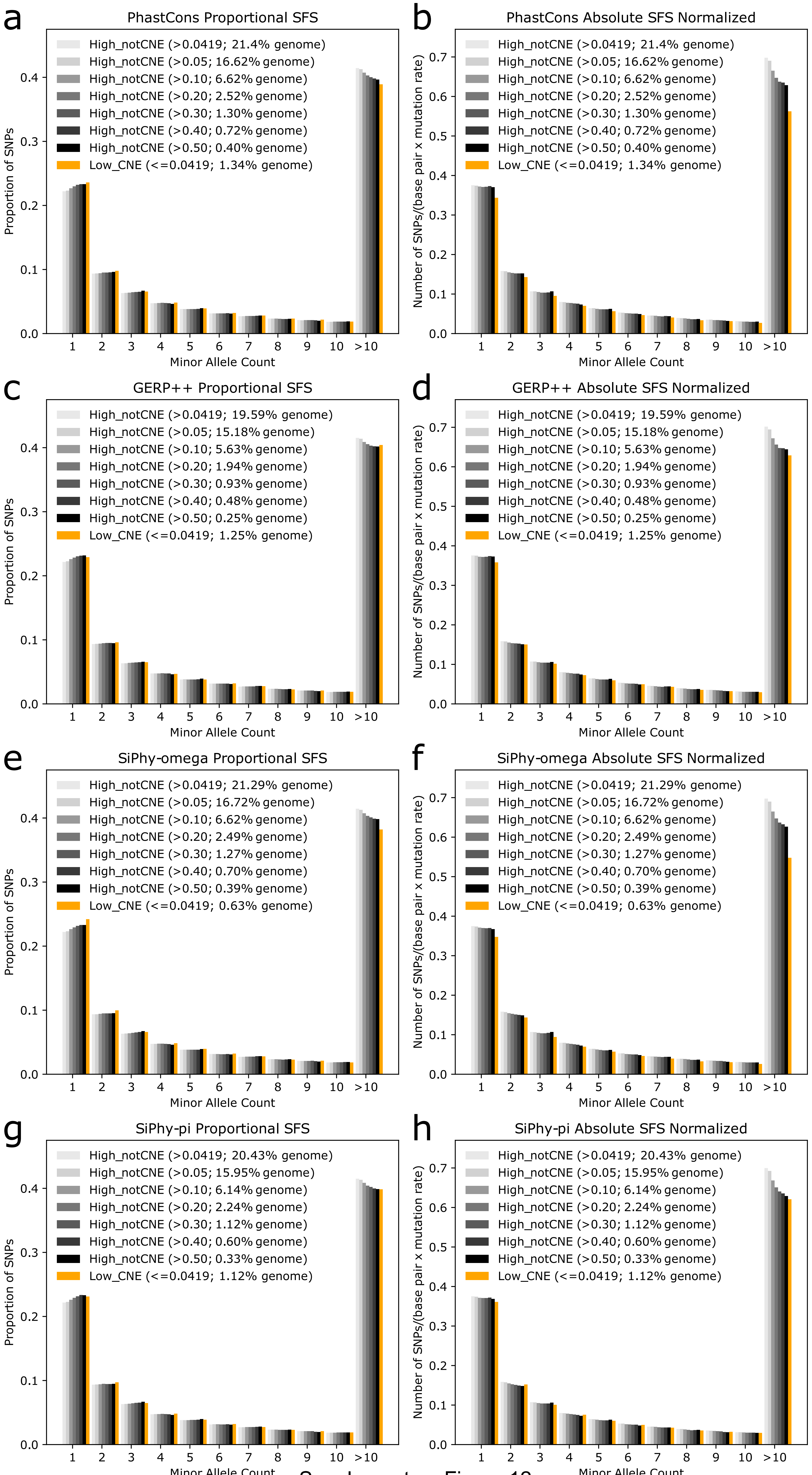


Supplementary Figure 12

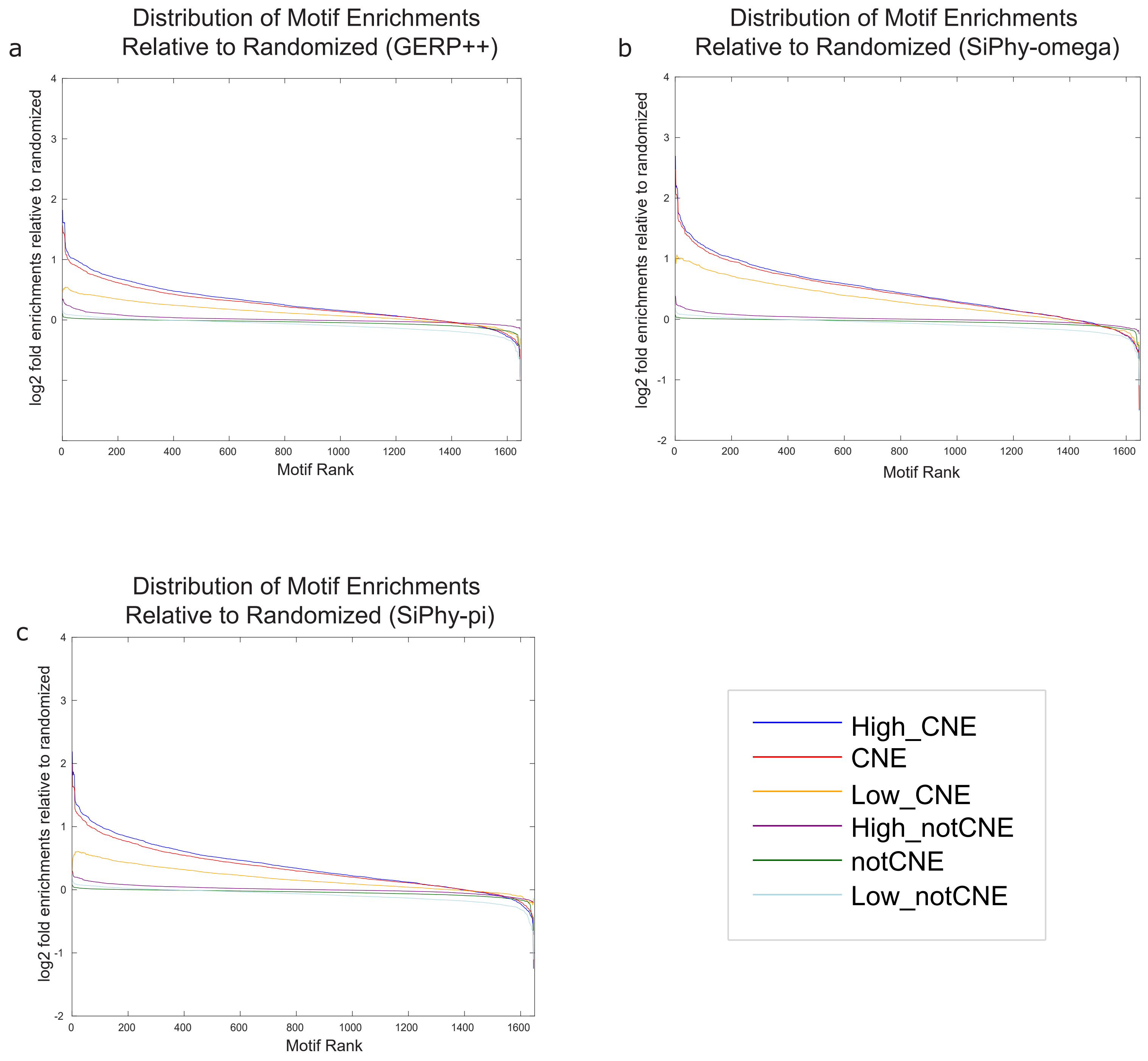


Supplementary Figure 13

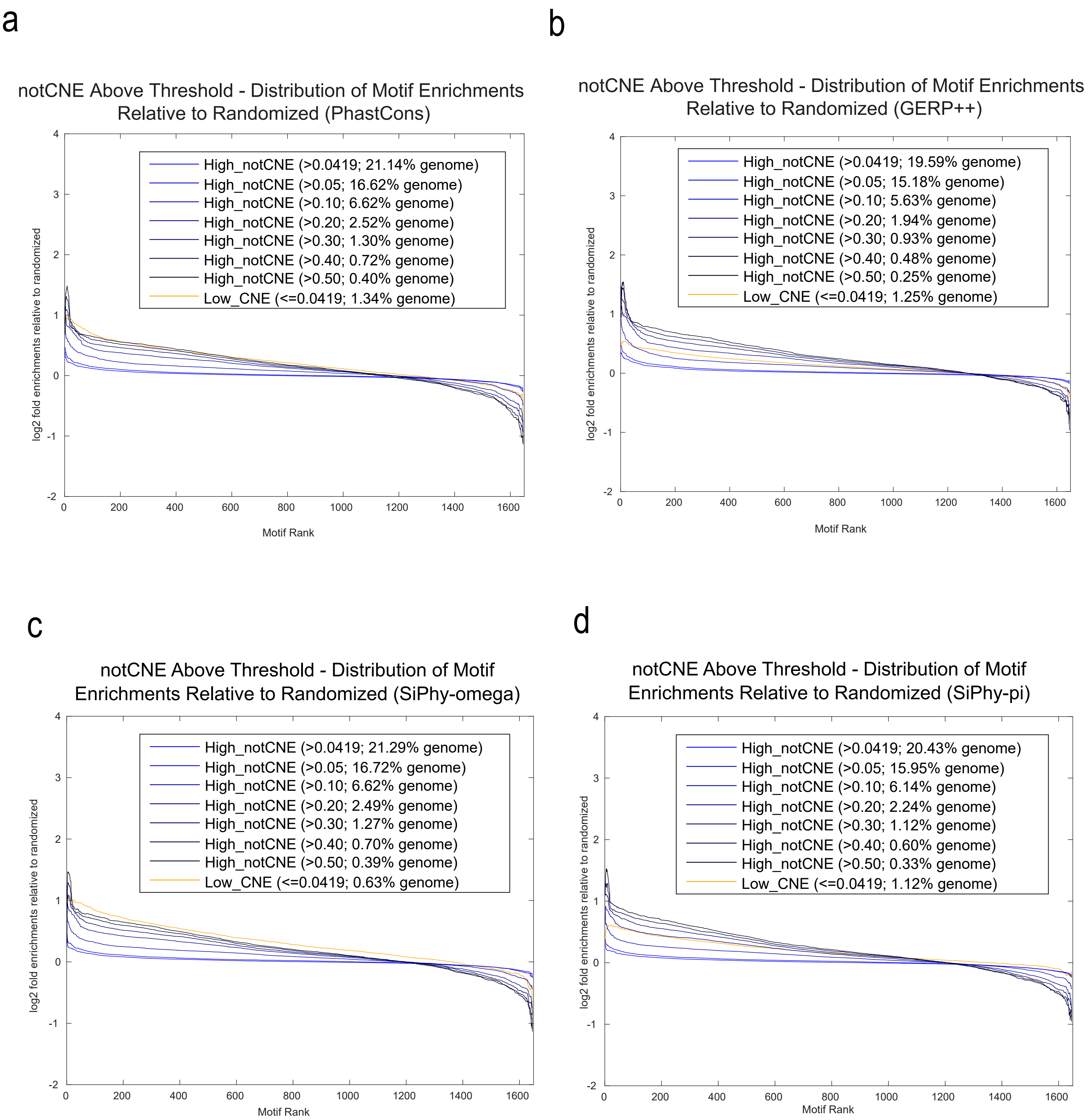


Supplementary Figure 14

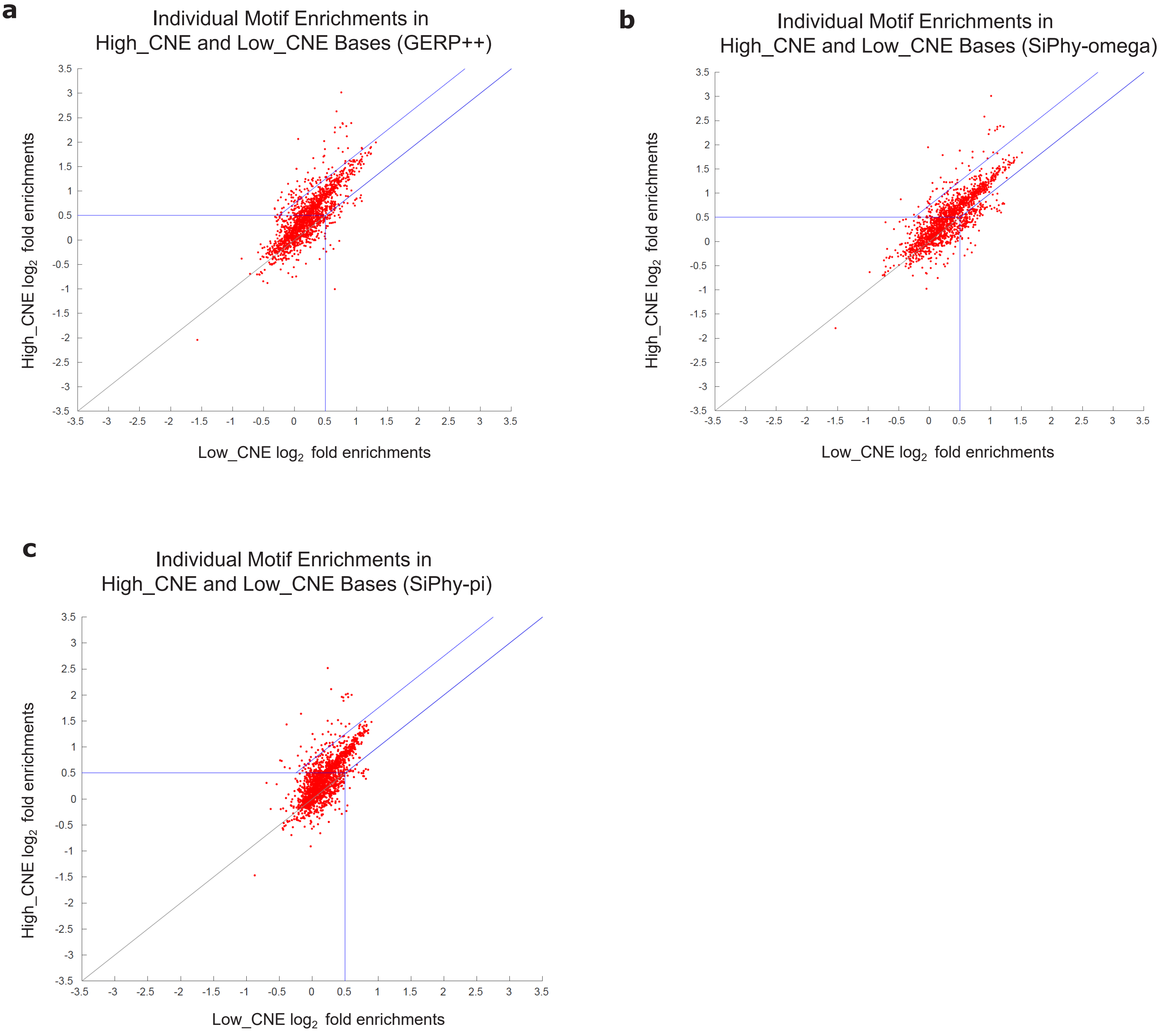


Supplementary Figure 15

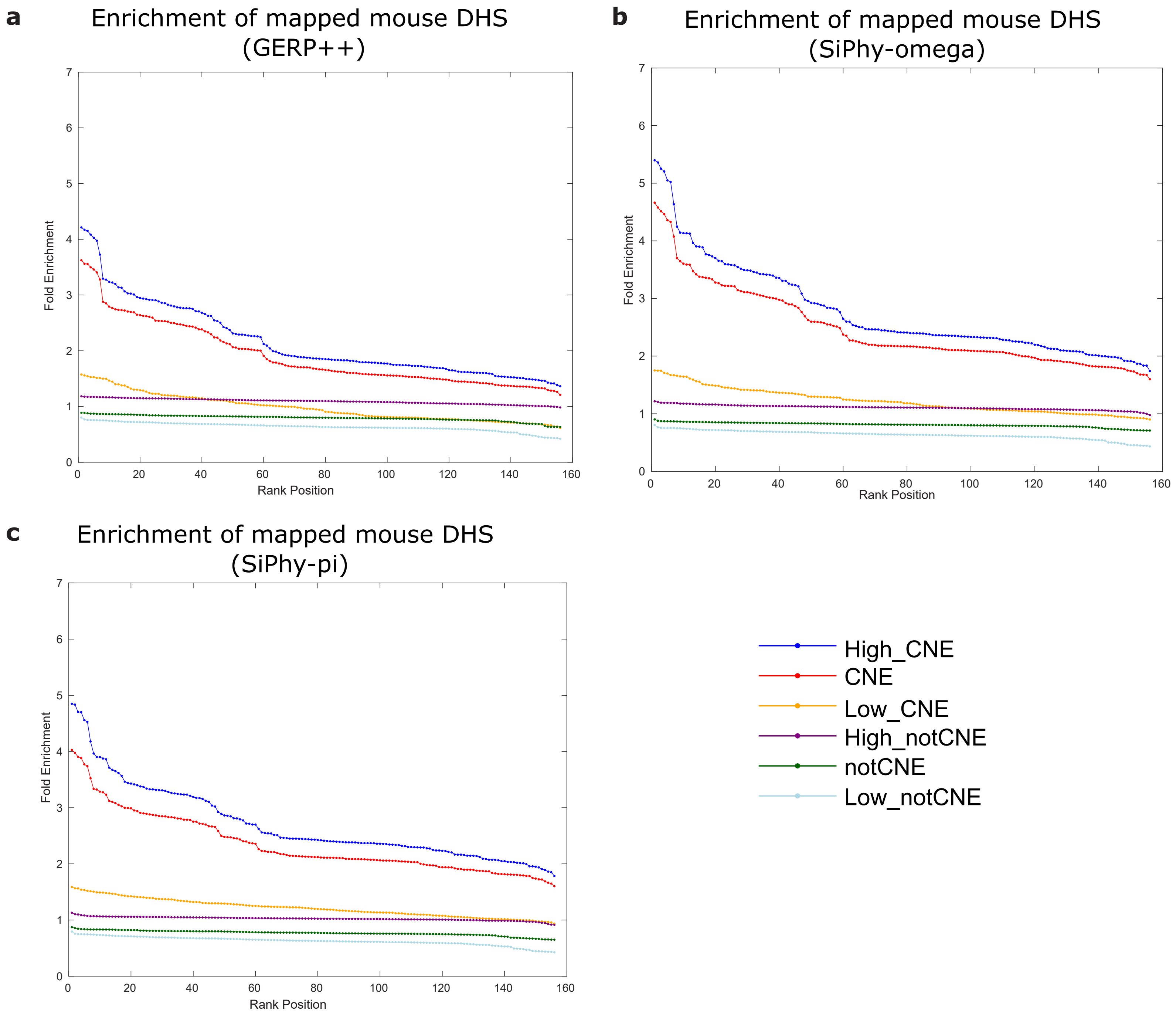


Supplementary Figure 16

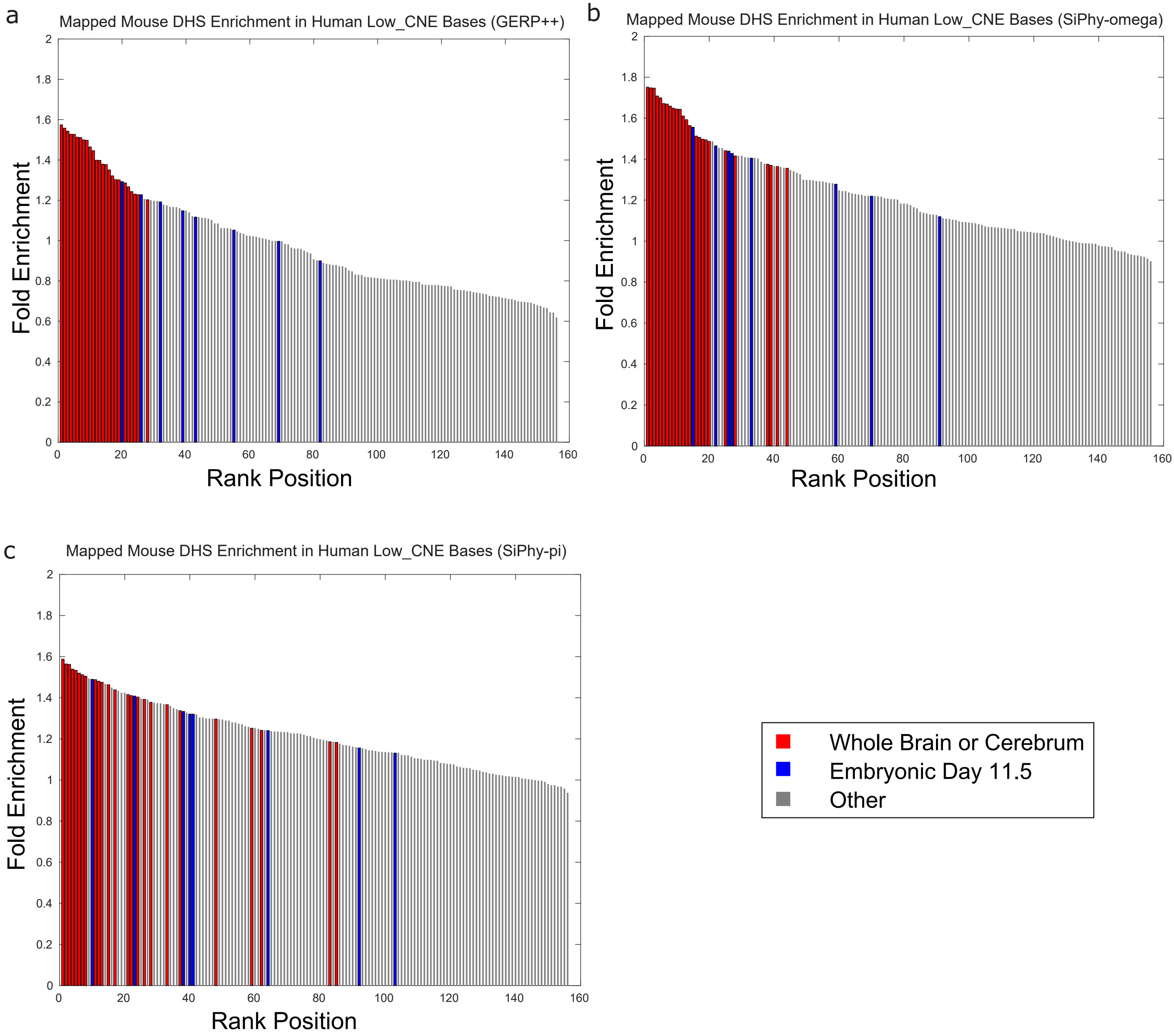


Supplementary Figure 17

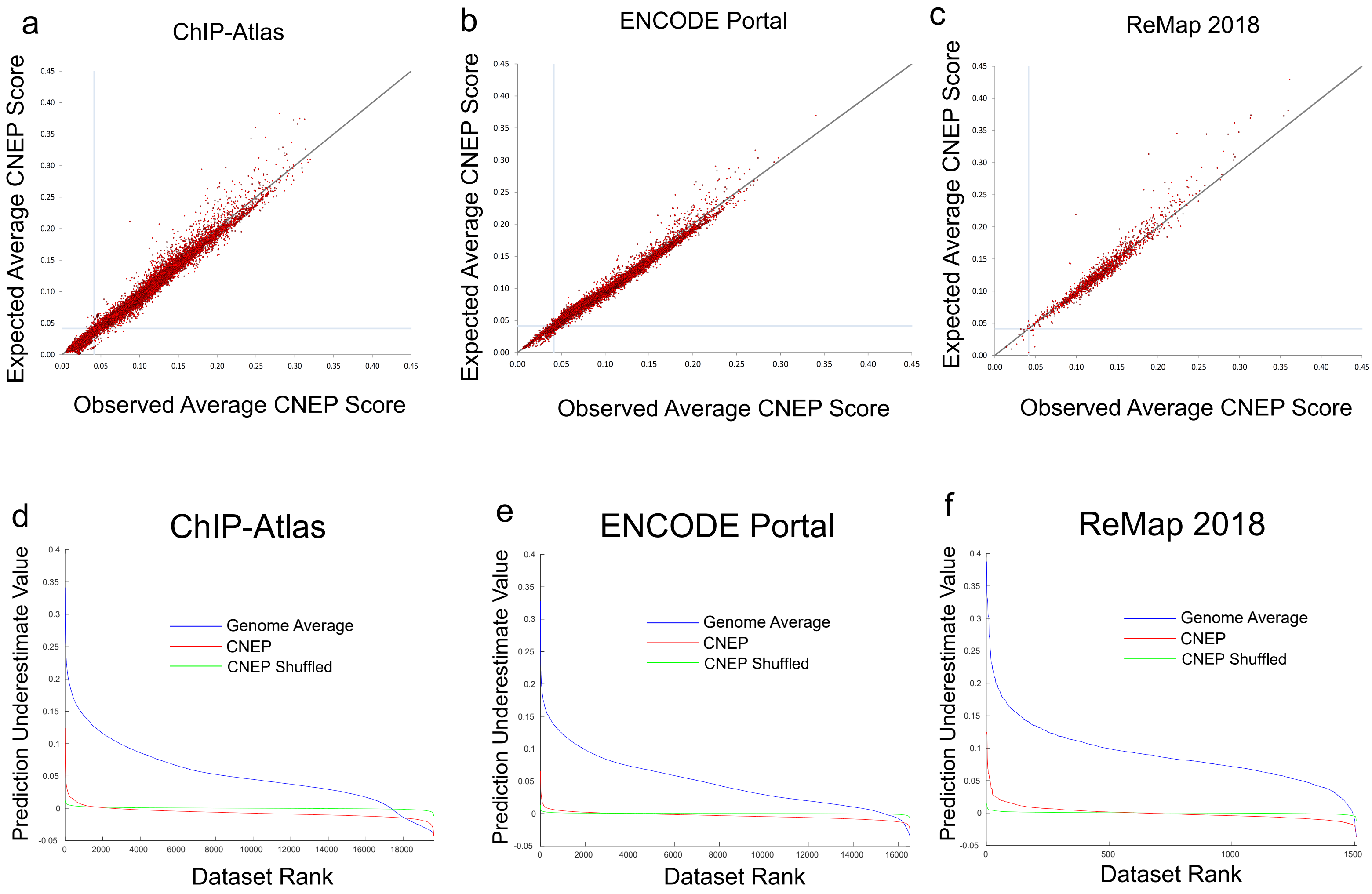


Supplementary Figure 18

a

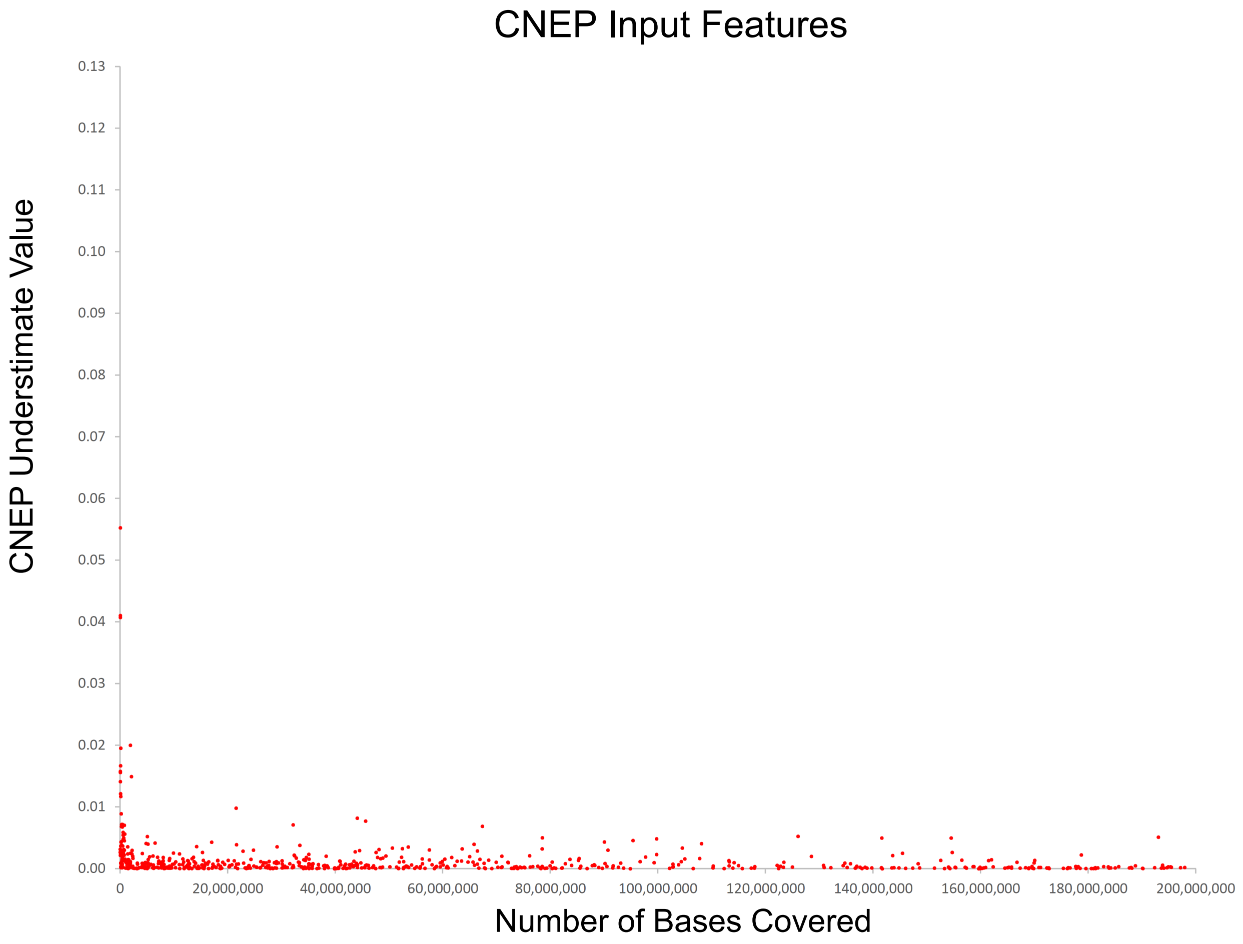


b

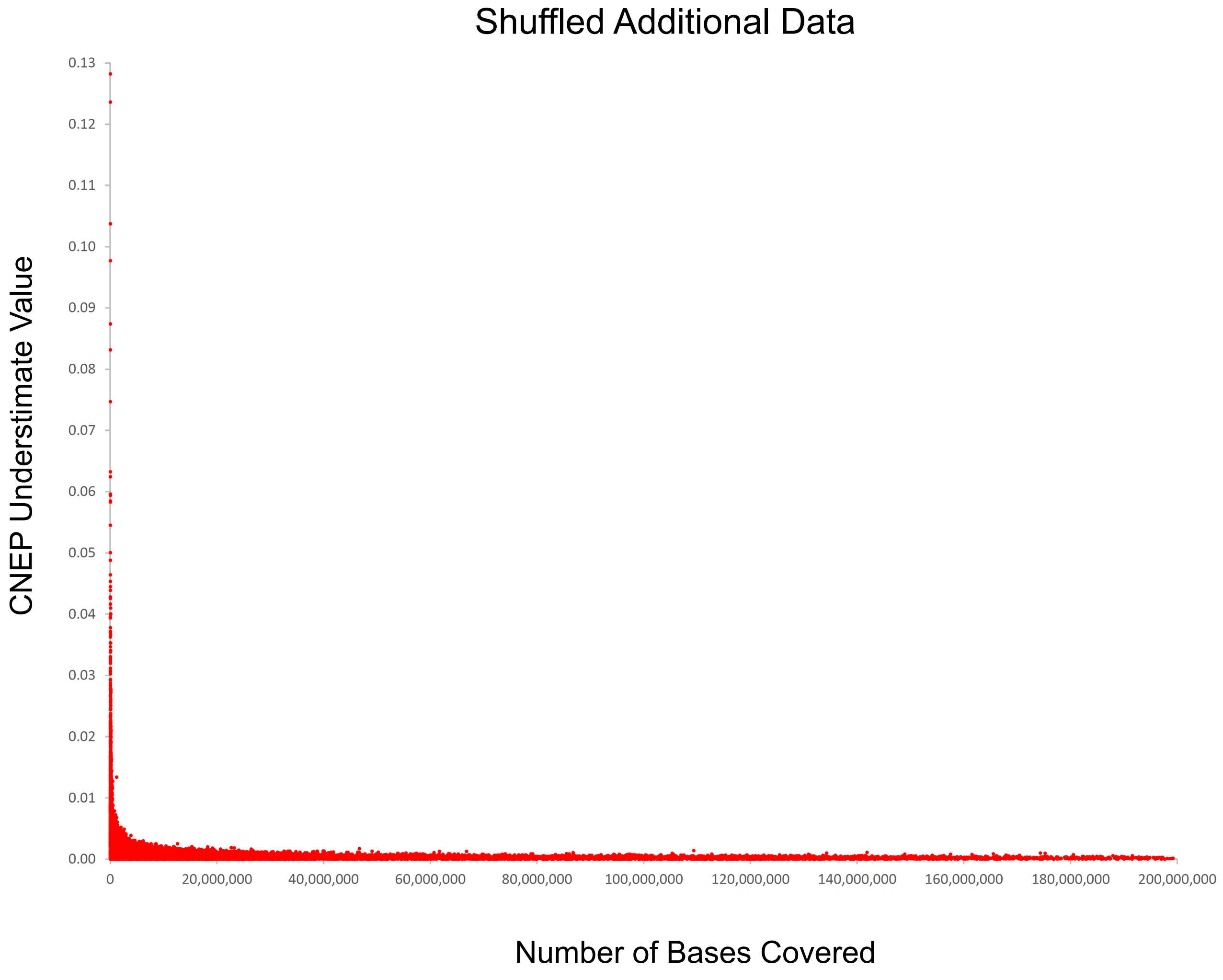


Supplementary Figure 19
