## Supplementary Legends for "Identification and characterization of constrained non-exonic bases lacking predictive epigenomic and transcription factor binding annotations"

### Supplementary Figure Legends

**Supplementary Figure 1: CNEP score correlation with scores based on individual constrained element sets.** The figure shows the genome-wide pairwise correlations between the predictions based on applying the CNEP method to each constrained element individually and the CNEP score, which was determined based on averaging predictions based on training on each of the four constrained element sets separately.

**Supplementary Figure 2: Cumulative distribution of CNEP scores for additional constrained element sets and genome coverage.** Similar to **Fig. 2a**, the graph shows the cumulative distribution of the CNEP score genome-wide (green), in constrained non-exonic (CNE) bases (red), and bases that are not in constrained elements and also not in exons (notCNE) (blue) for **(a)** GERP++, **(b)** SiPhy-omega and **(c)** SiPhy-pi. **(d)** The percent of genome covered by High_CNE, Low_CNE, CNE, High_notCNE, Low_notCNE, and notCNE bases. **(e)** Schematic illustration of the relationship between High_CNE, Low_CNE, CNE, High_notCNE, Low_notCNE, notCNE bases, and the CNEP score. The High_CNE, Low_CNE, High_notCNE, and Low_notCNE only contain non-exonic bases and all non-exonic bases are in one of those four sets.

**Supplementary Figure 3: Precision-recall and receiver operating characteristic curves for additional constrained element sets. (a-c)** The precision recall curves for the CNEP score for predicting non-exonic bases in constrained elements called by **(a)** GERP++, **(b)** SiPhy-omega, **(c)** SiPhy-pi. **(d-f)** The receiver operating characteristic (ROC) curves for the CNEP score for predicting non-exonic bases in constrained elements called by **(d)** GERP++, **(e)** SiPhy-omega, **(f)** SiPhy-pi. The areas under these ROC curves were 0.86, 0.86, and 0.84 respectively.

**Supplementary Figure 4: High_notCNE bases enrichment near CNE bases for additional constrained element sets. (a)** The plot shows the cumulative fraction of PhastCons High_notCNE bases at each distance to the nearest CNE base up to 3,000 bp. **(b)** The plot shows the fold enrichment for the cumulative number of PhastCons High_notCNE bases being within each distance to the nearest CNE base, up to 3,000bp. The same plots for other constrained element sets: **(c,d)** GERP++, **(e,f)** SiPhy-omega, and **(g,h)** SiPhy-pi.

**Supplementary Figure 5: Low_CNE bases enrichment near exons for additional constrained element sets. (a)** The plot shows the cumulative fraction of PhastCons Low_CNE bases at each distance to the nearest exon, up to 3,000bp. **(b)** The plot shows the fold enrichment for the cumulative number of PhastCons Low_CNE bases being within each distance to the nearest exon, up to 3,000bp. The same plots for other constrained element sets: **(c,d)** GERP++, **(e,f)** SiPhy-omega, and **(g,h)** SiPhy-pi.

**Supplementary Figure 6: Conservation state AUC and enrichments for additional constrained element sets.** This is an extended version of **Fig. 3a,b** showing the AUC, enrichment values, and average CSS-CNEP score in CNE bases for GERP++, SiPhy-omega, SiPhy-pi in addition to PhastCons.

**Supplementary Figure 7: Prediction of CNE bases for additional constrained element sets. (a)** ROC curves for the CNEP score identifying PhastCons CNE bases in specific ConsHMM conservation states. Curves are colored based on their corresponding state, as shown in **Fig. 3a**. **(b)** Same plot as in (a) except for GERP++. **(c)** Similar plots as shown in **Fig. 3c** except for GERP++ instead of PhastCons. **(d-g)** Similar plots as (c,d) except for **(d,e)** SiPhy-omega, and **(f,g)** SiPhy-pi.

**Supplementary Figure 8: Prediction of CNE bases more than 200bp from exons for additional constrained element sets.** Similar plots as shown in **Supplementary Fig. 7a** and **Fig. 3c** except the exon definition is first extended 200bp on each side so that only bases in constrained non-exonic elements more than 200bp from an exon are considered positives. Shown here for all four constrained element sets: **(a,b)** PhastCons, **(c,d)** GERP++, **(e,f)** SiPhy-omega, and **(g,h)** SiPhy-pi. State 1 now becomes the state with the highest AUC overall.

**Supplementary Figure 9: Prediction of Low_CNE bases among CNE bases.** Similar plots as shown in **Fig. 3d** except for **(a)** GERP++, **(b)** SiPhy-omega, and **(c)** SiPhy-pi.

**Supplementary Figure 10: Proportional site frequency spectrum (SFS) and absolute SFS normalized by average mutation rate for additional constrained element sets.** Similar plots as shown in **Fig. 4a,b** except for additional constrained element sets: **(a,b)** GERP++, **(c,d)** SiPhy-omega, and **(e,f)** SiPhy-pi.

**Supplementary Figure 11: Proportional SFS and absolute SFS normalized by average mutation rate controlling for background selection.** Similar plots as shown in **Fig. 4a,b** and **Supplementary Fig. 10** except reweighting positions to control for differences in estimated background selection. For that the proportional SFS, the weighting is such that the *B*-value distribution of variants in a set matches the *B*-value distribution of all variants. For the absolute density SFS normalized by mutation rate, the weighting is such that the *B*-value distributions of all considered non-exonic positions in a set matches the *B*-value distribution of all considered non-exonic positions. The plots are based on **(a,b)** PhastCons**, (c,d)** GERP++, **(e,f)** SiPhy-omega, and **(g,h)** SiPhy-pi constrained elements.

**Supplementary Figure 12: Proportional SFS and absolute SFS for subsets of High_notCNE bases at more stringent CNEP thresholds.** Similar plots as shown in **Fig. 4a,b** and **Supplementary Fig. 10** except comparing Low_CNE bases, which all have a CNEP score <=0.0419, High_notCNE bases, which all have a CNEP score >0.0419, and those High_notCNE bases with CNEP scores greater than 0.05, 0.10, 0.20, 0.30, 0.40, and 0.50 based on **(a,b)** PhastCons**, (c,d)** GERP++, **(e,f)** SiPhy-omega, and **(g,h)** SiPhy-pi constrained elements. The legend indicates the percent of the genome in each of the sets.

**Supplementary Figure 13: Distribution of motif enrichments for additional constrained element sets.** Similar plots as shown in **Fig. 4c** except for additional constrained element sets: **(a)** GERP++, **(b)** SiPhy-omega, and **(c)** SiPhy-pi.

**Supplementary Figure 14: Distribution of motif enrichments for subsets of High_notCNE bases at more stringent CNEP thresholds.** Similar plots to **Fig. 4c** and **Supplementary Fig. S13.** The plot shows the difference of the distribution of motif enrichments relative to the distribution for a randomized set of the motifs for High_notCNE bases, which all have a CNEP score >0.0419, and the subsets that have CNEP scores greater than 0.05, 0.10, 0.20, 0.30, 0.40, and 0.50 as indicated in the color legend. Also shown for comparison are the results for the Low_CNE bases, which all have a CNEP score <=0.0419. The x-axis is the rank position of the motif among the 1,646 motifs and the y-axis is the difference between the $\log_2$ fold enrichment based on the actual motif calls and the median of three randomized versions at the same rank position (**Methods**), shown for **(a)** PhastCons, **(b)** GERP++, **(c)** SiPhy-omega, and **(d)** SiPhy-pi constrained elements. The legend indicates the percent of the genome in each of the sets.

**Supplementary Figure 15: Scatter plot of individual motif enrichments for additional constrained element sets.** Similar plots as shown in **Fig. 4d** except for additional constrained element sets: **(a)** GERP++, **(b)** SiPhy-omega, and **(c)** SiPhy-pi. The blue lines separate the three regions used for the GO enrichment analysis, High_CNE strongly preferred, High_CNE moderately preferred, and Low_CNE preferred, where at least one of the Low_CNE or the High_CNE $\log_2$ enrichment is greater than or equal to 0.5 (**Supplementary Table 6**).

**Supplementary Figure 16: Distribution of enrichments for DNase I Hypersensitive Sites (DHS) from mouse for additional constrained element sets.** Similar plots as shown in **Fig. 4e** except for additional constrained element sets: **(a)** GERP++, **(b)** SiPhy-omega, and **(c)** SiPhy-pi.

**Supplementary Figure 17: Enrichments of DNase I Hypersensitive Sites (DHS) from mouse for additional constrained element sets.** Similar plots as shown in **Fig. 4f** except for additional constrained element sets: **(a)** GERP++, **(b)** SiPhy-omega, and **(c)** SiPhy-pi.

**Supplementary Figure 18: Expected vs. observed average CNEP scores for additional human datasets.** Similar scatter plots as shown in **Fig. 2b** except for **(a)** ChIP-atlas, **(b)** ENCODE portal, **(c)** and ReMap 2018 datasets using the retrospective CNEP score based on a subset of features available in 2015 (**Methods**). Each point corresponding to one dataset of peak calls. The x-axis shows the average CNEP score in bases covered by a peak, while the y-axis shows the expected CNEP score based on the peaks overlap with constrained non-exonic bases. Only datasets that cover at least 200kb are shown. The full table corresponding to these values can be found in **Supplementary Table 8**. The diagonal line is the y=x line. The vertical line corresponds to the genome-wide observed average CNEP score. The horizontal line corresponds to the genome-wide expected average CNEP score. **(d-f)** Plots for **(d)** ChIP-atlas **(e)** ENCODE portal, and **(f)** ReMap 2018 showing the distribution of prediction underestimate values for datasets with peaks covering at least 200kb. The prediction underestimate value for a dataset is the average difference between the expected CNEP score based on a subset of features available in 2015 (**Methods**) and the prediction value for each base covered by a peak. Results are shown for prediction values based on the genome-wide average expected CNEP score (blue) and the CNEP score (red). Also shown is the distribution of using the CNEP score for the prediction values, but applied to a shuffled version of each dataset (green). There was a relatively large gap in distribution based on using the observed CNEP score instead of genome-wide expected average CNEP score when computing the difference, highlighting that the CNEP score captures a relatively large amount of information about CNE bases. There is a difference in distributions between using the CNEP score on the actual peaks and a shuffled version of the peaks, though it was much smaller. These results suggest that CNEP captures most of the marginal information contained in any peak call dataset about the expectation on the frequency of CNE bases.

However, there are some datasets that capture some additional marginal information on CNE bases than given by the CNEP score.

**Supplementary Figure 19: CNEP underestimates values for input features and shuffled versions in retrospective analysis of additional data.** Similar plots to **Fig. 5** showing a scatter plot of CNEP underestimate values and bases covered for **(a)** the input features to CNEP and **(b)** shuffled versions of the additional datasets considered in **Fig. 5** based on a subset of features available in 2015 (**Methods**). The data in **Fig. 5** had greater underestimation values for the same genomic coverage than in both these controls, demonstrating additional marginal additive information about CNE bases in the additional datasets.

### Supplementary Table Legends

**Supplementary Table 1: Sources of features –** This table summarizes the sources of the epigenomics and TF binding data used as features to CNEP. Each feature corresponds to one file matching the URL up to the wild card (‘*’) except as indicated for the chromatin state and ReMap data. The last column indicates if the set of features were held out from training in the retrospective analysis.

**Supplementary Table 2: Observed and Expected average feature score for input features** – This table reports the observed average CNEP score for all 63,741 binary features provided to CNEP in bases in which the feature is positive. The table also reports the expected average CNEP score for the feature, which is the genome-wide frequency of constrained non-exonic elements overlapping bases in which the feature is present, averaged over the four different constrained element sets considered. The last column reports the number of bases in the genome for which the feature was positive. The features are listed in decreasing order of the observed average CNEP score.

**Supplementary Table 3: AUC performance at predicting constrained elements –** The first row of the table provides the AUC for the ROC curve for the CNEP score. The next row of the table shows the AUC based on just the score trained on that element set opposed to the average from the four elements, which led to similar performance. The next line shows the performance of CNEP based on a subset of features available by 2015 in the retrospective analysis. Below that is the baseline of counting the number of present features overlapping a base, counting the number of present features from DNase I experiments and just Roadmap Epigenomics DNase I experiments. Also shown is the performance of two existing scores, the Segway Encyclopedia 'conservation-associated activity score'(Libbrecht et al., 2019) and FitCons2 'cell-type integrated scores'(Gulko and Siepel, 2019). The AUC values are shown for each of the four constrained non-exonic element sets considered both when bases overlapping exons are included in the negative set and when they are excluded from both the positive and negative sets.

**Supplementary Table 4: Base set definition –** This table provides the definition used for CNE, Low_CNE, High_CNE, notCNE, Low_notCNE, High_notNCE bases for a given constrained element set.

**Supplementary Table 5: Motif enrichments –** This table reports the $\log_2$ fold enrichment for 1,646 motifs from (Kheradpour and Kellis, 2014) for bases in Low_CNE and High_CNE for each of the four constrained element sets considered. Motifs are ordered based on the enrichment in Low_CNE bases for PhastCons.

**Supplementary Table 6: TF Motif GO enrichments –** For each constrained element set, there is a separate sheet with three tables reporting GO enrichments for TF subsets corresponding to three motif subsets. The subsets, in order, correspond to High_CNE strongly preferred, High_CNE moderately preferred, and Low_CNE motif subsets (**Methods**). The columns are the ID of the GO category, GO category name, uncorrected p-value, corrected p-value, and fold enrichment. Only GO enrichments that had a corrected p-value <=0.10 are shown.

**Supplementary Table 7: Mouse DHS enrichments for human Low_CNE bases** – This table reports the fold enrichment of mouse DHS mapped to human in Low_CNE bases relative to a randomized version of the DHS. Included in the table are enrichments for the four constrained element sets considered for each of the 156 mouse DNase I hypersensitivity experiments.

**Supplementary Table 8: Retrospective analysis of information in additional human datasets –** This table reports separately for each dataset included in the ChIP-atlas, ENCODE portal, and ReMap 2018 compendia as well as for GENCODE exons present in v28 and not v19: (i) observed average CNEP score, (ii) expected average CNEP score, (iii) number of base pairs covered by the dataset, (iv) difference between the expected and observed average CNEP score. For these analyses the CNEP score was derived based on 10,836 features available as of 2015. Additionally, the antigen, cell type, and cell type class from the database provided metadata is also reported for ChIP-atlas. The assay, biosample, and where applicable experiment target from the metadata are reported for the ENCODE portal data. Information about the dataset is included in the dataset ID for ReMap.

**Supplementary Table 9: ChIP-atlas cell type class enrichments** – The table reports the top enrichments for cell type class in terms of p-value for the 209 ChIP-atlas datasets that had a CNEP underestimate value greater than 0.02 restricting to those datasets whose peaks covered at least 200kb. The columns are the name of the cell type class, uncorrected p-value, corrected p-value, and fold enrichment.
